# Profiling of the drug resistance of thousands of Src tyrosine kinase mutants uncovers a regulatory network that couples autoinhibition to catalytic domain dynamics

**DOI:** 10.1101/2021.12.05.471322

**Authors:** Sujata Chakraborty, Ethan Ahler, Jessica J. Simon, Linglan Fang, Zachary E. Potter, Katherine A. Sitko, Jason J. Stephany, Miklos Guttman, Douglas M. Fowler, Dustin J. Maly

## Abstract

Kinase inhibitors are effective cancer therapies but resistance often limits clinical efficacy. Despite the cataloguing of numerous resistance mutations, our understanding of kinase inhibitor resistance is still incomplete. Here, we comprehensively profiled the resistance of ∼3500 Src tyrosine kinase mutants to four different ATP-competitive inhibitors. We found that ATP-competitive inhibitor resistance mutations are distributed throughout Src’s catalytic domain. In addition to inhibitor contact residues, residues that participate in regulating Src’s phosphotransferase activity were prone to the development of resistance. Unexpectedly, we found that a resistance-prone cluster of residues located on the top face of the N-terminal lobe of Src’s catalytic domain contributes to autoinhibition by reducing catalytic domain dynamics, and mutations in this cluster led to resistance by lowering inhibitor affinity and promoting kinase hyperactivation. Together, our studies demonstrate how drug resistance profiling can be used to define potential resistance pathways and uncover new mechanisms of kinase regulation.

## INTRODUCTION

Small molecule, ATP-competitive kinase inhibitors have revolutionized the treatment of specific cancers (Cohen et al., 2021; Gharwan and Groninger, 2016; Zhang et al., 2009). Unfortunately, as with many targeted cancer therapies, the efficacy of ATP-competitive kinase inhibitors is frequently limited by the rapid development of resistance. Mechanisms of ATP-competitive kinase inhibitor resistance are often complex and heterogeneous (Garraway and Janne, 2012; Lovly and Shaw, 2014) but point mutations that render a target kinase less sensitive to inhibition are common (Krishnamurty and Maly, 2010). How do these kinase mutations result in resistance to an ATP-competitive inhibitor? One frequently observed mechanism is through the disruption of favorable binding interactions with the inhibitor. For example, mutation of the “gatekeeper” residue, which sits near the back of the ATP-binding pocket, to a bulkier residue that restricts the size of an adjacent hydrophobic pocket is one of the major pathways of resistance observed in the clinic for several kinases (Gibbons et al., 2012; Patel et al., 2020). Similarly, other inhibitor contact residues are also commonly observed sites of ATP-competitive inhibitor resistance (Balzano et al., 2011). However, the mechanistic basis for many clinically observed mutations that do not directly influence drug binding is less clear, and even mutations that affect inhibitor binding can confer resistance through additional mechanisms. For example, while gatekeeper residue mutations often lead to a reduction in inhibitor affinity they can also decrease inhibitor efficacy by increasing the ability of ATP to compete for ATP-binding site occupancy (Yun et al., 2008) and have been shown to lead to hyperactivation of some tyrosine kinases (Azam et al., 2008). Therefore, studies that allow the comprehensive profiling of ATP-competitive inhibitor resistance are valuable in identifying potential sites of inhibitor resistance and also providing new insight into kinase regulation and activity.

To obtain insight into different mechanisms of drug resistance, we used a deep mutational scanning (DMS) approach (Fowler et al., 2010; Fowler and Fields, 2014) to investigate the impact of nearly every mutation in Src kinase’s catalytic domain on the efficacy of a panel of ATP-competitive inhibitors. Using this approach, we first established Src’s pattern of resistance to the clinically approved inhibitor dasatinib. As expected, mutations at sites within the catalytic domain that interact with dasatinib were capable of conferring resistance. We also observed that residues involved in the autoinhibition of Src’s phosphotransferase activity were particularly prone to the development of resistance to dasatinib. Next, we profiled a matched set of conformation-selective, ATP-competitive inhibitors to compare how resistance emerges in response to different modes of ATP-competitive inhibition. We identified a number of unique resistance mutations for each conformation-selective inhibitor and a shared set of residues that were highly susceptible to the development of resistance. Further investigation of a spatially defined cluster of residues located on the solvent-exposed N-terminal lobe of Src’s catalytic domain, which we refer to as the β1/β2 resistance cluster, demonstrated that mutations at these residues were capable of reducing ATP-competitive inhibitor affinity. Biochemical analyses revealed that many mutations in the β1/β2 resistance cluster led to hyperactivation of Src’s phosphotransferase activity and that this structural element also contributes to the autoinhibited state of Src by reducing the dynamics of the catalytic domain’s N-terminal lobe, which resistance mutations release. Together, our results show how a previously unrecognized component of Src’s autoinhibitory regulatory network is a major site for the potential development of drug resistance.

## RESULTS

### A yeast-based growth assay for profiling drug resistance

To comprehensively profile mechanisms of inhibitor resistance in Src, we used an assay that relies on the correlation between Src’s phosphotransferase activity and its toxicity in *S. cerevisiae* (Ahler et al., 2019; Brugge et al., 1987; Kritzer et al., 2018). We reasoned that it would be possible to determine the drug sensitivity of individual Src variants by measuring Src-mediated yeast toxicity in the presence of various ATP-competitive inhibitors (**Figure 1A**). Prior to performing a parallel analysis of Src mutants, we validated that ATP-competitive inhibitors, including the highly potent Src inhibitor dasatinib (**Figure 1B**), were capable of dose-dependently rescuing the growth inhibition that resulted from expressing wild-type (WT), myristoylated, full-length Src (Src^myr^, see **Table S1** for all Src constructs) in yeast (**Figure S1A-D**). Next, we transformed a barcoded library (**Table S2**) of Src^myr^ in which the catalytic domain was mutagenized into yeast, performed outgrowth in the presence of 25 or 100 μM dasatinib and collected samples at various time points. At each time point, we extracted plasmid DNA, amplified barcodes, and then deeply sequenced the barcode amplicons to quantify the frequency of each Src variant. From these variant frequency data, we calculated activity scores for ∼3,500 single amino acid Src mutants at two different dasatinib concentrations by averaging two biological replicates (**Figure 1C-1F, S1E-F**).

**Figure 1.**
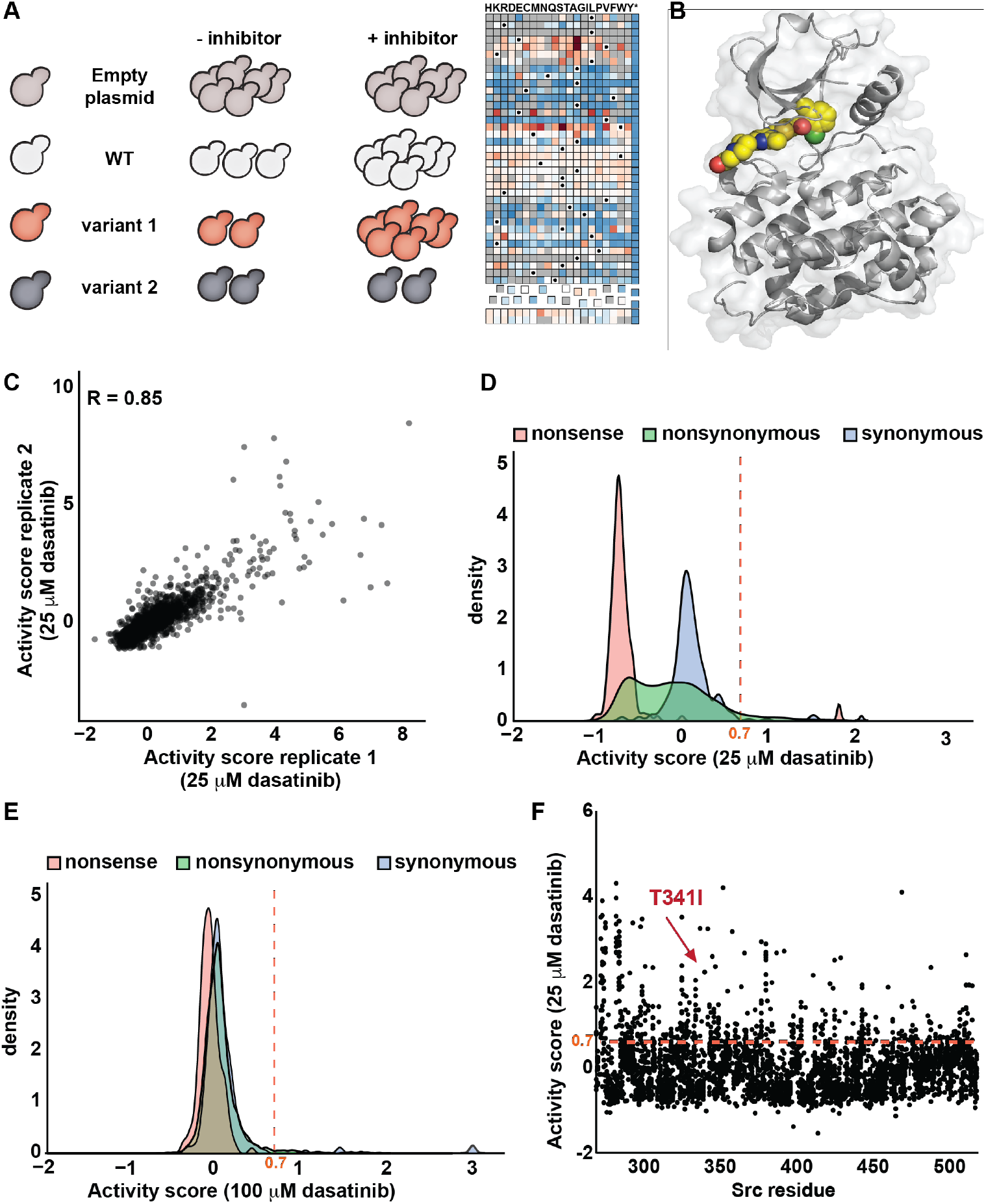
A yeast-based assay for the parallel analysis of drug resistance of Src tyrosine kinase mutants. (A) Schematic of the yeast-based growth assay used to quantify the drug resistance of thousands of Src mutants (B) Co-crystal structure of Src’s catalytic domain (PDB ID: 3G5D) bound to dasatinib (*colored spheres*). (C) Scatterplot showing the correlation (Pearson’s R = 0.85) between the activity scores obtained in 25 μM dasatinib-treated yeast for two independent transformations of the Src^myr^ variant library. (D, E) Activity score distributions of all synonymous and nonsynonymous (missense and nonsense) Src^myr^ variants in yeast treated with 25 μM (D) or 100 μM (E) dasatinib. The red dashed line indicates the activity score value (>2 standard deviations above the mean of the synonymous WT distribution) we defined as dasatinib resistant. (F) Scatterplot showing activity scores for every single amino acid substitution at every residue in Src’s catalytic domain for yeast transformed with the Src^myr^ library and treated with 25 μM dasatinib. The red dashed line indicates the activity score value we defined as dasatinib resistant, and each black dot represents the activity score derived from averaging two replicates. See also Table S1 and S2 and Figure S1.

For each concentration of dasatinib treatment, we defined a Src mutant as being drug resistant if it showed an activity score greater than two times the standard deviation of the mean of the synonymous distribution (**Figure 1D, 1E**). We found that patterns of dasatinib resistance looked similar for both dasatinib concentrations but with a narrower distribution of activity scores for 100 μM dasatinib treatment (**Figure 1D, 1E**). Therefore, we further analyzed the data obtained from 25 μM dasatinib treatment, where a wider spectrum of mutational effects on resistance were apparent.

### Profiling of dasatinib resistance in Src

We assembled our large-scale mutagenesis data into a sequence-activity map covering more than 75% of all possible single mutants of Src’s catalytic domain (**Figure 2A, Table S3**). About 12% of these single mutants were defined as dasatinib-resistant based on our classification criteria.

**Figure 2.**
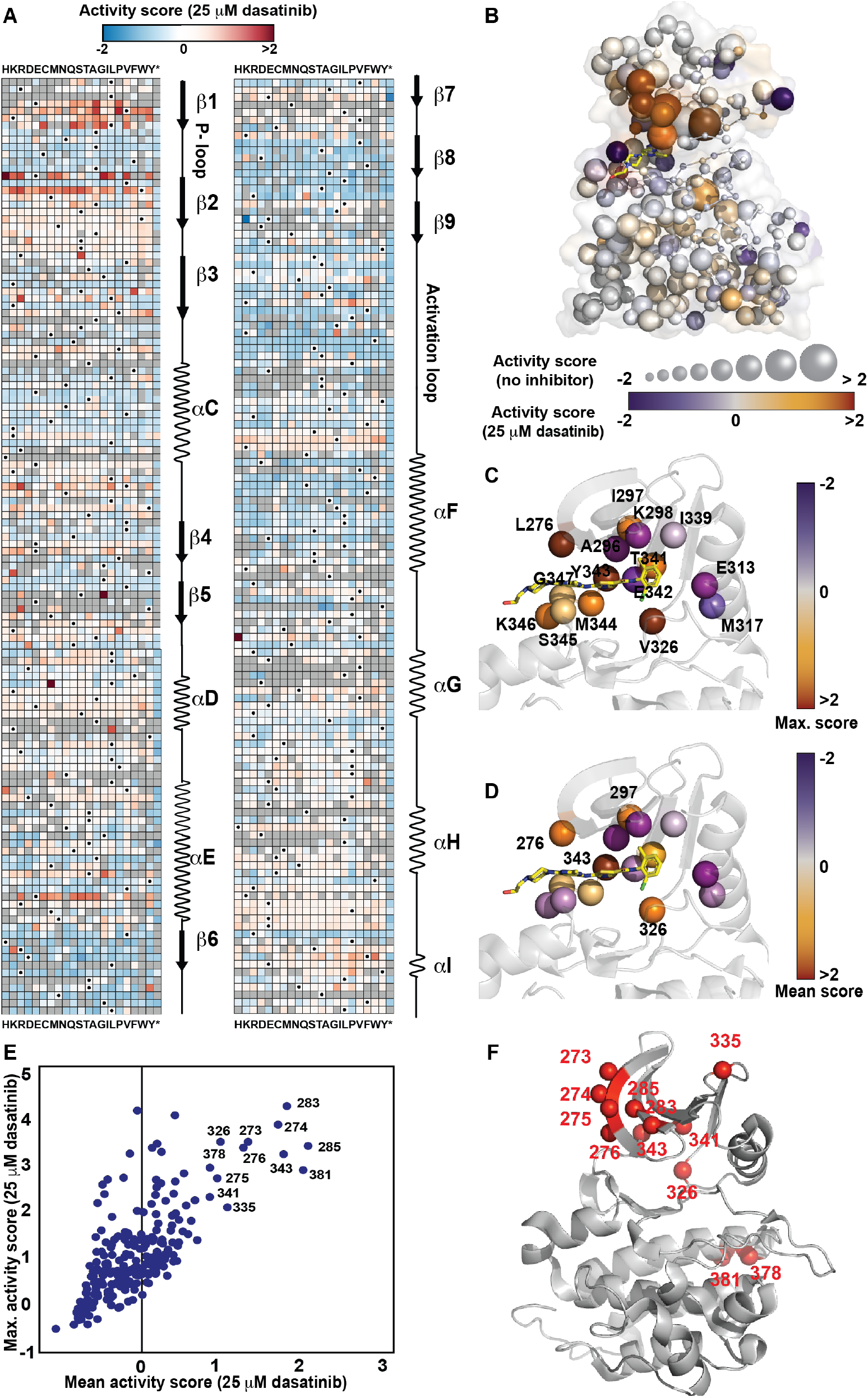
Mapping of dasatinib resistance mutations in Src. (A) Sequence-activity score map for all residues in Src’s catalytic domain for the Src^myr^ variant library treated with 25 μM dasatinib. Black dots represent the wild-type amino acid and gray tiles indicate missing data. Secondary structure and functional motif annotations were obtained from the ProKinO database. (B) Mean (position-averaged) activity scores for each residue in Src’s catalytic domain (PDB ID: 3G5D) for the Src^myr^ variant library in yeast treated with DMSO. Sphere size indicates position-averaged activity scores in the absence of dasatinib (Ahler *et al*., 2019); color indicates position-averaged activity scores in the presence of 25 μM dasatinib (*this work*). Nonsense mutants were excluded from the average score. (C) Co-crystal structure (PDB ID: 3G5D) of Src’s catalytic domain bound to dasatinib (*yellow sticks*) showing maximum activity scores in the presence of 25 μM dasatinib for all residues that interact with dasatinib. (D) Co-crystal structure (PDB ID: 3G5D) of Src’s catalytic domain bound to dasatinib (*yellow sticks*) showing mean activity scores in the presence of 25 μM dasatinib for all residues that interact with dasatinib. (E) Correlation plot of maximum activity scores and mean activity scores in the presence of 25 μM dasatinib for each residue in Src’s catalytic domain. (F) Crystal structure (PDB ID: 3G5D) of Src’s catalytic domain with the twelve residues that were defined as “resistance prone” shown as red spheres. See also Table S3 and Figure S2.

Consistent with the notion that preservation of phosphotransferase activity is required for a mutation to confer drug resistance, we observed that almost all dasatinib-resistant mutations were either classified as activating (∼65%) or WT-like (∼30%) in a comprehensive analysis of Src’s phosphotransferase activity that we previously performed (**Figure 2B, S2A**) (Ahler *et al*., 2019). While elevated phosphotransferase activity appears to be a common dasatinib resistance mechanism, activity scores obtained in the absence or presence of dasatinib were not strictly correlated (**Figure 2B, S2B**), suggesting that diverse mechanism of resistance were captured. Our sequence-activity map revealed that dasatinib-resistant mutations were distributed throughout the catalytic domain, especially in multiple residues in the β1 and β2 strands of the N-terminal lobe and the αD and αE helices of the C-lobe (**Figure 2A, 2B**). At many positions, only a few mutations conferred dasatinib resistance, while at a subset of residues most substitutions demonstrated dasatinib resistance (**Figure 2A**).

Mutations that perturb inhibitor contact residues in the ATP-binding pocket are a common mechanism of resistance to kinase inhibitors (Persky et al., 2020). We explored whether our large-scale mutagenesis data captured this mechanism of dasatinib resistance by mapping the maximum and mean (position-averaged) activity scores that were obtained in the presence of 25 µM dasatinib onto the fifteen residues in Src’s ATP-binding site that interact with dasatinib (**Figure 2C, 2D**). As expected, mutations at a number of residues that line the ATP-binding pocket led to substantial dasatinib resistance, including the well-characterized T341I gatekeeper mutation (Azam *et al*., 2008; Krishnamurty and Maly, 2010). In addition to the gatekeeper residue, residues Leu276 (located on the β1 strand), Val326 (located between the αC helix and β4 strand) and Tyr343 (located in the hinge region) all showed high mean activity scores in the presence of 25 µM dasatinib (**Figure 2D, 2E**) and appeared to be highly susceptible to the development of resistance. In total, at least one substitution led to resistance at two-thirds of the residues that interact with dasatinib, consistent with the ATP-binding pocket being a site of significant potential drug resistance. Our results are consistent with a recent screen of cyclin-dependent kinase 6 (CDK6)’s resistance to the ATP-competitive inhibitor palbociclib (Persky *et al*., 2020). Despite the slightly different binding poses of palbociclib and dasatinib within the ATP-binding pockets of CDK6 and Src, respectively, resistance mutations occurred at many of the same contact residues, and a similar percentage of contact residues were capable of acquiring inhibitor resistance (**Figure S2C, S2D**).

Our sequence-activity map shows that there are several residues where numerous substitutions led to dasatinib resistance, suggesting that they represent sites that are particularly prone to the development of drug resistance. To quantitatively identify resistance-prone sites, we determined all residues in Src’s catalytic domain that demonstrated mean (position-averaged) activity scores greater than our defined drug-resistance cutoff for an individual mutant in the presence of 25 µM dasatinib (**Figure 2E, 2F, S2E**). In total, twelve residues meet the “resistance-prone” definition. While four of the twelve resistance-prone residues (Leu276, Val326, Thr341, Tyr343) interact with dasatinib (**Figure 2C, 2D**), the remaining eight either possess sidechains that are directed away from the ATP-binding site or are distal to the site of dasatinib binding. We previously observed that all eight of the resistance-prone residues that do not interact with dasatinib contained a large number of activating mutations in the absence of dasatinib and are likely involved in the regulation of Src’s phosphotransferase activity (Ahler *et al*., 2019). Ala378 and Glu381 are components of the regulatory αF pocket in the C-terminal lobe that contributes to the autoinhibition of Src’s phosphotransferase activity through an interaction with the N-terminal SH4 domain. Resistance-prone residues Glu273, Val274, Lys275, Glu283, and Trp285 form a solvent-exposed, spatially defined cluster of residues located on the β1 and β2 strands of the N-terminal lobe and appear to contribute to Src’s autoinhibition by an unknown mechanism. Thus, elevated phosphotransferase activity appears to be a general mechanism for acquiring dasatinib resistance in Src.

### Src mutational resistance to different modes of ATP-competitive inhibition

We further explored the intersection between the regulation of Src’s phosphotransferase activity and drug resistance by determining how resistance arises in response to different modes of ATP-competitive inhibition (**Figure S3A**) (Fang et al., 2020; Potter et al., 2020; Ranjitkar et al., 2010). We profiled our Src variant library in the presence of each of a matched set of pyrazolopyrimidine-based ATP-competitive inhibitors that stabilize structurally distinct conformations of Src’s ATP-binding site (**Figure 3A**). Inhibitor **1** contains a 3-phenol at the C-3 position of the pyrazolopyrimidine scaffold that can potentially form a hydrogen-bond with the sidechain of Glu313 in the αC helix, leading to stabilization of the active conformation (αC helix-in; **Figure S3A**). Inhibitor **2** contains a pharmacophore at the C-3 position that promotes the outward rotation of the αC helix to an inactive conformation (αC helix-out). We also attempted to profile an inhibitor (inhibitor **3**) that contains a pharmacophore at the C-3 position of the pyrazolopyrimidine scaffold that stabilizes a flipped, inactive conformation of Src’s activation loop (DFG-out) but is otherwise identical to inhibitors **1** and **2**. However, inhibitor **3** did not achieve sufficiently high intracellular concentrations in yeast to inhibit WT Src^myr^. Therefore, we used inhibitor **4**, an analog of **3** that achieved high enough concentrations in yeast to inhibit WT Src^myr^ (**Figure S1D**), for our profiling of conformation-selective inhibitor resistance.

**Figure 3.**
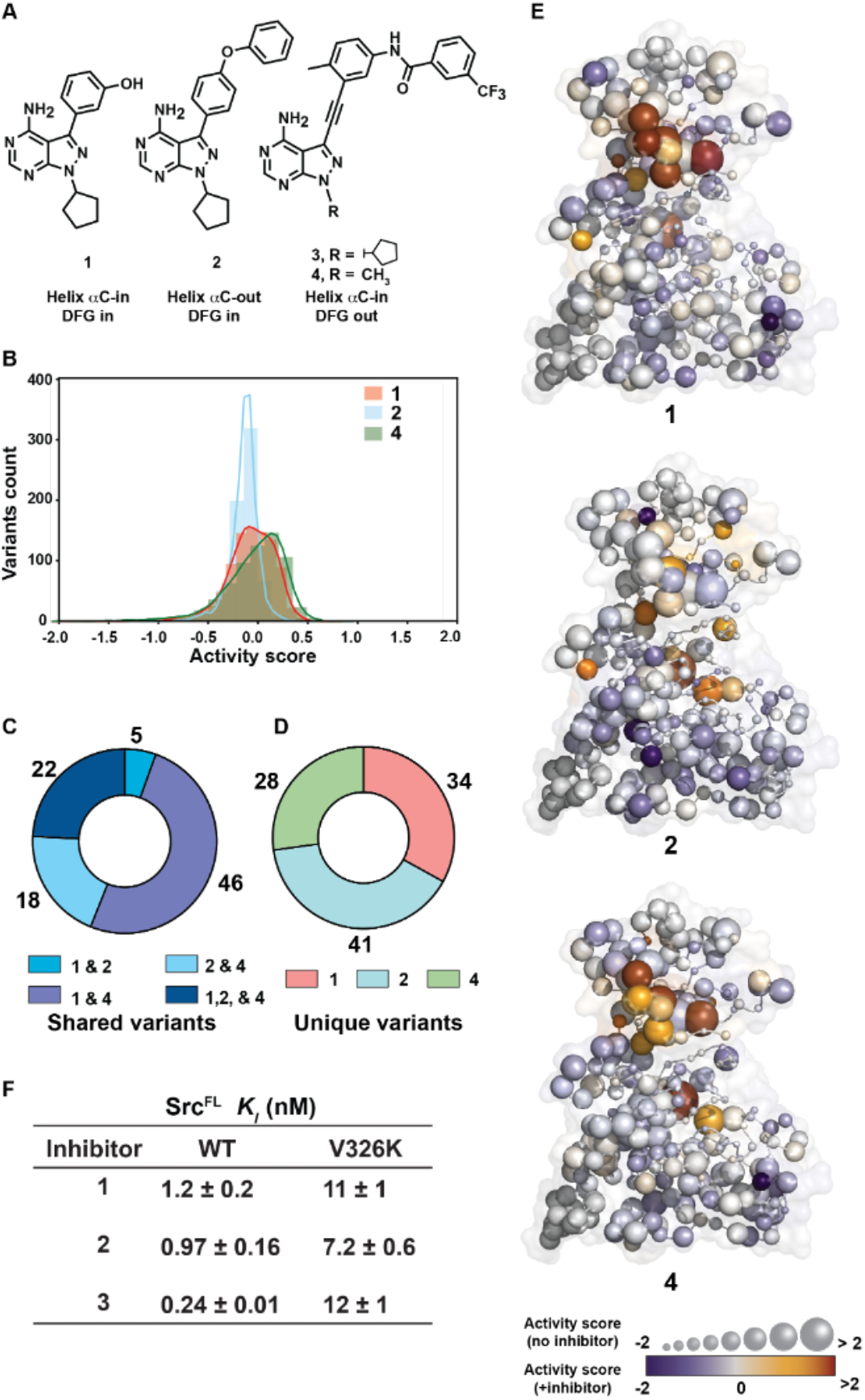
Parallel analysis of mutations conferring Src resistance to conformation-selective, ATP-competitive inhibitors. (A) Structures of the conformation-selective, ATP-competitive inhibitors that were profiled in our yeast-based growth assay. (B) Activity score distribution of all non-synonymous Src variants in yeast treated with inhibitors **1, 2** or **4**. (C) Donut plot showing the number of shared inhibitor-resistant mutants. (D) Donut plot showing the number of unique inhibitor-resistant mutants for inhibitors **1, 2** and **4**. (E) Mean activity scores for each residue in Src’s catalytic domain (PDB ID: 3G5D) for the Src^myr^ variant library treated with conformation-selective inhibitor **1** (*top*), **2** (*middle*) or **4** (*bottom*) and in absence of inhibitor (data from (Ahler *et al*., 2019)). Mean activity scores in the presence of each inhibitor are represented as color, while activity scores in the absence of inhibitors are represented by sphere size. (F) *K*_*I*_ values (n=3, mean ± SEM) of conformation-selective inhibitors **1**-**3** for WT and V326K Src^FL^. See also Table S4 and Figure S3.

After transforming the Src^myr^ variant library into yeast, we performed outgrowth in the presence of either inhibitor **1, 2**, or **4** and collected samples at various time points. For each inhibitor, we calculated normalized activity scores for ∼3,500 Src mutants (**Figure 3B, Table S4**). As for dasatinib, a Src variant was defined as being resistant if its activity score in the presence of a specific conformation-selective inhibitor was greater than two times the standard deviation of the mean of the synonymous distribution. Using this definition, we identified 107, 86, and 114 Src mutants that were resistant to inhibitors **1, 2**, and **4**, respectively (**Figure S3B**). Similar to Src’s resistance to dasatinib, almost all variants that were resistant to inhibitors **1, 2**, or **4** were either classified as activating or WT-like in our previous analysis of Src’s phosphotransferase activity (**Figure S3C**). In comparing the overlap between resistance mutations, we observed 22 mutations occurring at eight residues that were resistant to all three conformation-selective inhibitors (**Figure 3C, S3D**). Inhibitors **1** and **4** shared the highest overlap (∼75%) in resistance mutations, suggesting that their shared ability to stabilize an active conformation of Src’s αC helix makes them susceptible to similar mechanisms of resistance. Despite these similarities, we observed that ∼25-45% of all resistance mutations for a particular inhibitor were unique (**Figure 3D**), revealing resistance mechanisms that are likely specific to different modes of conformation-selective inhibition.

To provide a comprehensive overview of which regions are generally resistant to different modes of ATP-competitive inhibition, we mapped mean activity scores for each conformation-selective inhibitor onto the catalytic domain of Src (**Figure 3E**). Inhibitors **1** and **4** shared similar patterns of resistance, with the solvent-facing N-terminal lobe of the catalytic domain representing the main region of resistance. While residues in the same region of Src’s N-terminal lobe also demonstrated resistance to inhibitor **2**, unique resistance mutations to this αC helix-out-stabilizing inhibitor were also observed through the entirety of Src’s kinase domain. In particular, three residues (Leu300, Ile337, and Leu413) contained multiple mutations that uniquely conferred strong resistance to inhibitor **2**. Leu300 is located on the linker that connects the αC helix to the β-sheet of the N-terminal lobe and Ile337 is located on the face of the β4 strand that is directed towards helix αC. A plausible explanation for how mutations at these residues confer resistance is that they negatively influence the ability of helix αC to adopt the inactive “out” conformation that is required for inhibitor **2** to be accommodated in Src’s ATP-binding pocket (Chakraborty et al., 2019).

We observed six residues (Leu276, Trp285, Val326, Thr341, Tyr343, and Glu381) that fulfill our definition of being resistance-prone to all three conformation-selective inhibitors (**Figure 3E, S3D**). These six residues were also classified as being resistance-prone for dasatinib (**Figure 2F**). The sidechains of Leu276, Val326, Thr341, and Tyr343 are all directed towards Src’s ATP-binding pocket and mutations at these residues most likely confer resistance by perturbing inhibitor contacts. For example, Thr341 is commonly referred to as the gatekeeper residue and many substitutions at this residue have been shown to lead to inhibitor resistance by restricting the size of the ATP-binding pocket. We speculated that mutations at Val326 (**Figure S3E)**, which contains a sidechain that is directed towards the adenine ring of ATP, provided resistance through a similar mechanism. To test this prediction, we performed phosphotransferase activity and inhibition assays with WT and an inhibitor-resistant variant (V326K) of purified full-length Src (Src^FL^). We observed that purified V326K Src^FL^’s K_M_ for ATP (K_M_[ATP]) and phosphotransferase activity were very similar to WT Src^FL^’s (**Figure S3F**). Consistent with the V326K mutation leading to reduced inhibitor affinity, purified V326K Src^FL^ displayed K_I_ values for inhibitors **1, 2** and **3** that were 5- to 50-fold higher than WT Src^FL^ (**Figure 3F**). Interestingly, the tyrosine kinases BCR-Abl, Alk, and c-KIT also contain a valine at an equivalent position in their ATP-binding sites and drug-resistant mutations have been observed at this position in the clinic (**Figure S3G**). Thus, this region of the ATP-binding site appears to be a site of inhibitor resistance for a number of tyrosine kinases.

### Characterization of a resistance-prone cluster of residues in the N-terminal lobe of Src’s catalytic domain

Six residues (Glu273, Val274, Lys275, Leu276, Glu283, and Trp285) that possess multiple drug-resistant mutants form a spatially defined cluster on the top face of the β1 and β2 stands in the N-terminal lobe of Src’s catalytic domain (**Figure 4A, S4A-E**). Despite the high number of mutations that conferred resistance at the six residues within this cluster, which we hereafter refer to as the β1/β2 resistance cluster, the sidechains of all but Leu276 are solvent exposed and directed away from the site of ATP binding in Src (**Figure S4A**). Therefore, we were curious why this cluster of solvent-exposed residues is so prone to the development of drug resistance. To determine whether mutations within the β1/β2 resistance cluster confer resistance by reducing Src’s affinity for ATP-competitive inhibitors, we determined the K_I_ values of inhibitors **1**-**3** for purified Src^FL^ constructs containing drug-resistant E283M or W285T mutations (**Figure 4B**). We found that the presence of either the E283M or W285T mutation led to increased K_I_ values for inhibitors **1**-**3** relative to WT Src^FL^ but had a negligible effect on the K_M_ for ATP (**Figure 4B, S4G**). Furthermore, we found that both mutations conferred similar levels of inhibitor resistance. Therefore, mutations in the β1/β2 resistance cluster confer resistance, in part, by lowering the affinity of inhibitors for Src’s ATP-binding site.

**Figure 4.**
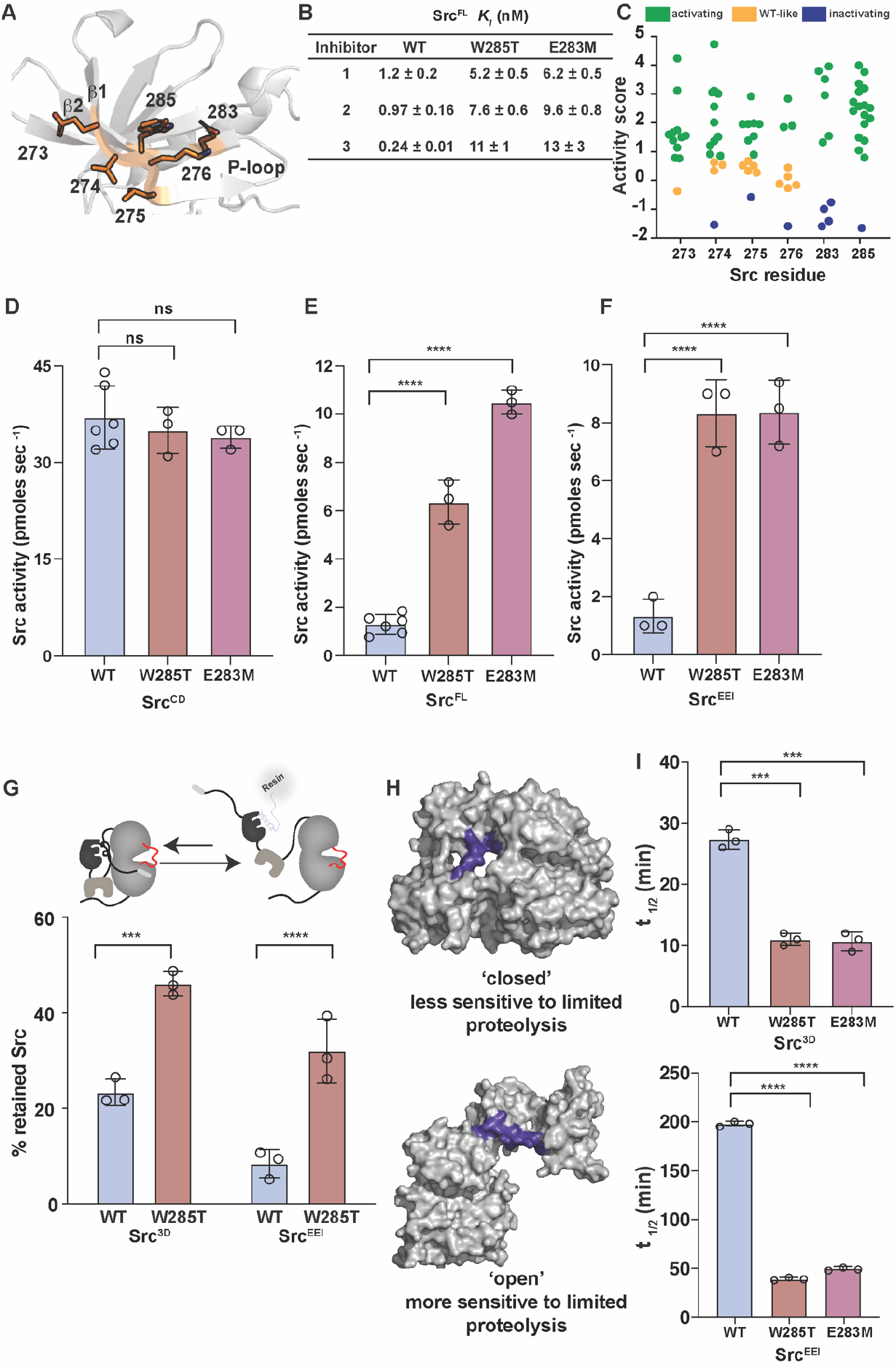
Biochemical characterization of the β1/2 resistance cluster. (A) Top-down view of the N-terminal lobe of Src’s catalytic domain (PDB ID: 1Y57) with the sidechains of all six residues in the β1/2 resistance cluster shown in orange. (B) *K*_*I*_ values of inhibitors **1**-**3** for the WT, E283M and W285T mutants of Src^FL^ (n=3, mean ± SEM,). (C) Activity scores for every substitution at each residue in the β1/2 resistance cluster in the absence of an inhibitor. Values were determined in a previous study (Ahler *et al*., 2019). (D) Phosphotransferase activity of purified WT, W285T, or E283M Src^CD^ (n=3-5, ns = non-significant, P > 0.05). (E) Phosphotransferase activity of purified WT, W285T, or E283M Src^FL^ (n=3-5, \*\*\*\**P < 0*.*0001*). (F) Phosphotransferase activity of purified WT, W285T or E283M Src^EEI^ (n=3, \*\*\*\**P < 0*.*0001*). (G) Schematic of the SH3 domain pull-down assay (*top*). To detect global conformation, Src constructs were incubated with an immobilized SH3 domain ligand. Closed, SH3 domain-engaged Src is unable to bind to the resin, whereas open, SH3 domain-disengaged Src binds. After washing, retained Src is eluted and quantified by western blot analysis (bottom). Percent retained WT or W285T Src^3D^ (left, n=3, *\*\*\*P < 0*.*001*) and WT or W285T Src^EEI^ (right, n=3, \*\*\*\**P < 0*.*0001*) in the SH3 domain pull-down assay. (H) Crystal structure (PDB ID: 2SRC) of Src^3D^ in the closed (*top*) and open (bottom) global conformations. The SH2-catalytic domain linker (violet) of the open conformation of Src^3D^ is more sensitive to proteolysis by thermolysin than the closed global conformation. (I) Half-life values of purified WT, E283M, or W285T Src^3D^ (*top*, n=3, \*\*\**P < 0*.*001*) and WT, E283M, or W285T Src^EEI^ (*bottom*, n=3, \*\*\*\**P < 0*.*0001*) in the thermolysin limited proteolysis assay. See also Figure S4.

### Residues in the β1/β2 resistance cluster modulate autoinhibition of Src

We previously observed that all six residues within the β1/β2 resistance cluster also contained multiple activating mutations in the absence of inhibitors (**Figure 4C**), suggesting that this region may serve a role in modulating Src’s phosphotransferase activity (Ahler *et al*., 2019). Therefore, we next determined how activating mutations in the β1/β2 resistance cluster influence Src’s phosphotransferase activity with purified Src constructs **(Figure S4F**). Interestingly, we found that the E283M and W285T mutations had a negligible effect on the phosphotransferase activity of a construct (Src^CD^) consisting solely of the catalytic domain of Src (**Figure 4D, S4G**). However, we observed that both mutations were activating in the context of Src^FL^ (**Figure 4E**), suggesting that activating mutations in the β1/β2 resistance cluster release regulatory domain-mediated autoinhibition. Consistent with the β1/β2 resistance cluster influencing regulation mediated by Src’s SH2 and SH3 domains, and not the N-terminal SH4 and unique domains, the E283M and W285T mutations showed a similar level of activation relative to WT in a Src^3D^ construct, Src^EEI^, that possesses SH2 domain interaction-enhancing mutations in the C-terminal tail that promote a similar level of autoinhibition as Src^FL^ (**Figure 4F**). Together, our data support a model wherein residues in the β1/β2 resistance cluster reinforce the autoinhibition provided by the SH2 and SH3 domain regulatory apparatus, which activating mutations release.

We speculated that activating mutations in the β1/β2 resistance cluster increase Src’s phosphotransferase activity by reducing levels of intramolecular SH2 and SH3 domain regulatory engagement. To test this prediction, we assessed how mutations affect intramolecular regulatory domain engagement levels with two biochemical assays. First, we measured intramolecular SH3 domain engagement levels with an immobilized SH3 domain ligand pull-down assay (**Figure 4G**). Consistent with activating β1/β2 resistance cluster mutations leading to increased phosphotransferase activity by reducing the autoinhibitory engagement of Src’s regulatory apparatus, W285T Src^3D^’s association with an immobilized SH3 domain ligand was more than 2-fold greater than WT Src^3D^’s (**Figure 4G, S4H**). Furthermore, the W285T mutation dramatically increased the ability of the more regulatory domain-engaged Src^EEI^ construct to intermolecularly interact with the immobilized SH3 domain ligand relative to WT Src^EEI^ (**Figure 4G, S4H**). We next used the rate of proteolysis of the flexible linker that connects Src’s SH2 domain to its CD (SH2-CD linker) by the metalloprotease thermolysin to characterize how β1/β2 resistance cluster mutations affect intramolecular regulatory domain engagement. Previous studies have demonstrated an inverse correlation between the rate of thermolysin cleavage, measured as intact protein half-life (t_1/2_), of the SH2-CD linker and intramolecular SH2 and SH3 regulatory domain engagement levels (**Figure 4H**) (Agius et al., 2019; Fang *et al*., 2020). Concordant with the SH3 domain pull-down results, the SH2-CD linkers of E283M and W285T Src^3D^ were proteolyzed 2-3 times more rapidly than WT Src^3D^ **(Figure 4I, S4I**). The E283M and W285T mutations also dramatically increased the rate of proteolytic cleavage of Src^EEI^’s SH2-CD linker (**Figure 4I, S4J**). Thus, residues in the β1/β2 resistance cluster appear to influence the level of intramolecular engagement of Src’s SH2 and SH3 domains.

### The W285T mutation promotes a more dynamic N-terminal lobe of Src

While our biochemical analyses suggest that activating mutations in the β1/β2 resistance cluster increase Src’s phosphotransferase activity by promoting a more open, regulatory domain-disengaged global conformation, the mechanistic basis for this effect was unclear. To the best of our knowledge, no previous structural or biochemical studies have suggested that residues within the β1/β2 resistance cluster participate in regulatory interactions. Furthermore, residues in the β1/β2 resistance cluster are separated by >10 Å from the nearest regulatory interface of autoinhibited Src (**Figure 5A, 5B**). Thus, we performed Hydrogen-Deuterium eXchange Mass Spectrometry (HDX-MS) on Src^3D^ to provide an unbiased analysis of how activating β1/β2 resistance cluster mutations affect the global conformational dynamics of Src (Boczek et al., 2019; Hochrein et al., 2006; Potter *et al*., 2020). Specifically, we compared the deuterium backbone exchange kinetics of W285T Src^3D^ relative to WT Src^3D^. We subjected identical concentrations of WT Src^3D^ and W285T Src^3D^ to standard D_2_O exchange conditions and samples were quenched and processed at various time points using established methods. Using this protocol, we were able to monitor the exchange kinetics of peptic peptides covering ∼85% of WT Src^3D^’s and W285T Src^3D^’s sequence (**Table S5**).

**Figure 5.**
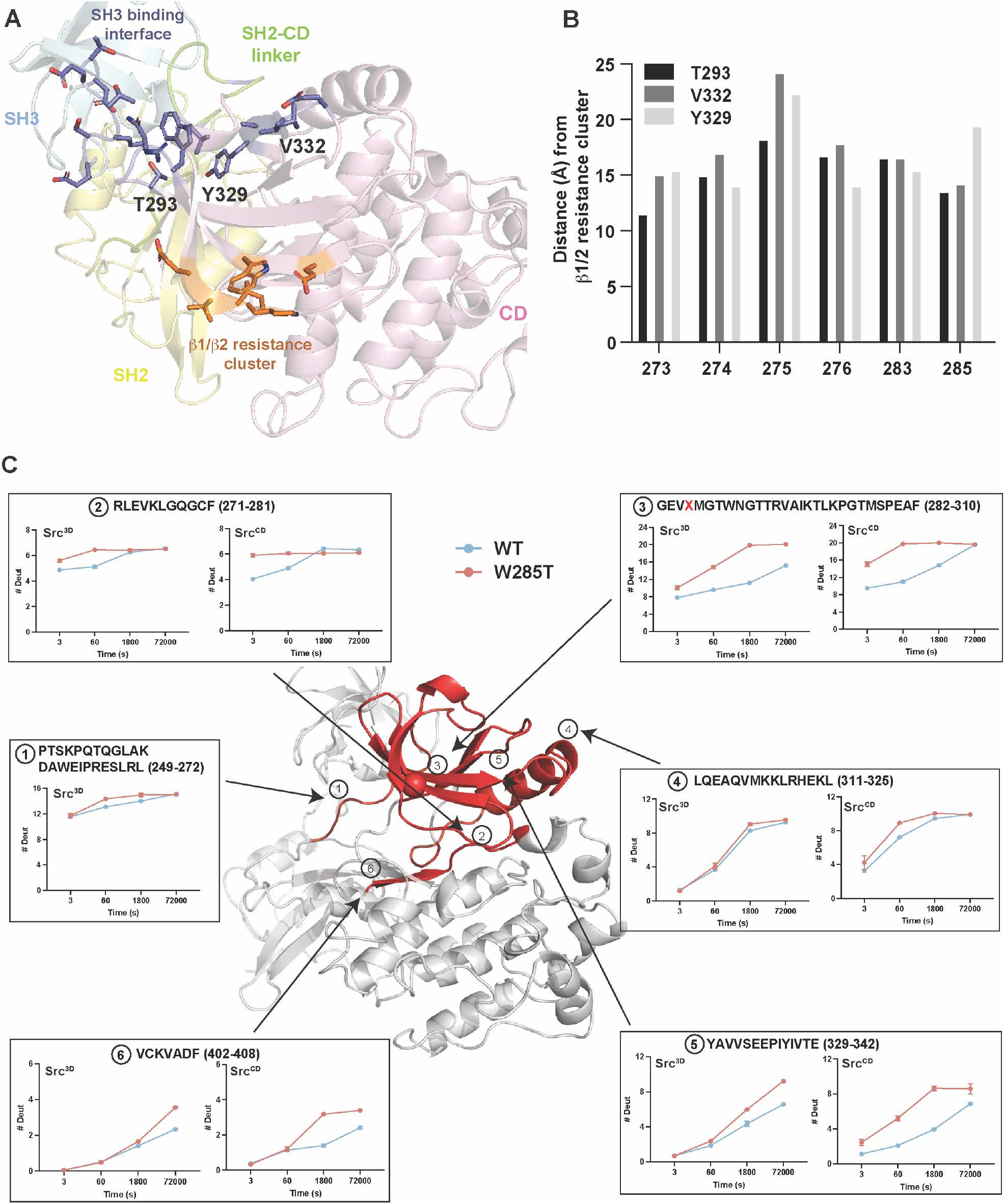
Conformational dynamics of W285T Src. (A) Structure of autoinhibited Src^3D^ (PDB ID: 2SRC) showing the sidechains of residues in the β1/β2 resistance cluster (*orange*) and the sidechains of residues in the nearest regulatory interface (*blue*, SH3 domain interface) in Src’s catalytic domain. (B) Plot showing the distances between the β-carbons of the three residues (T293, Y329, and V332) in the SH3 domain regulatory interface closest to the β1/2 resistant cluster and the β-carbons of residues in the β1/2 resistant cluster. (C) HDX-MS analysis of purified Src^3D^ (WT compared to W285T) and Src^CD^ (WT compared to W285T). Deuteration differences between WT and W285T Src^3D^ and Src^CD^ are plotted on the crystal structure of Src^3D^ (PDB ID: 2SRC). Regions that show increased deuterium uptake (significant differences in exchange were assessed using the hybrid significance threshold at 99% CI as well as consistency among all observed overlapping peptides, Hageman et al., 2019) in W285T Src^3D^ and/or Src^CD^ relative to WT Src^3D^ and/or Src^CD^, respectively, are shown in red, while regions that show no difference are shown in white. The values shown at each timepoint represent the mean ± SEM (n=3). See also Table S5.

Mapping differences in deuterium exchange kinetics between W285T Src^3D^ and WT Src^3D^ onto a structure of autoinhibited Src^3D^ (**Figure 5C**) revealed that a large portion of Src’s catalytic domain in W285T Src^3D^ underwent faster exchange kinetics than in WT Src^3D^. Consistent with β1/β2 resistance cluster mutations leading to a reduction in the intramolecular engagement of Src’s regulatory SH2 and SH3 domain apparatus (**Figures 4G, 4I**), peptic peptides covering the SH2-catalytic domain linker (peptide 1: 249-272) and the SH3 domain interface (peptide 3: 282-310) of autoinhibited Src demonstrated increased solvent accessibility in W285T Src^3D^ relative to WT Src^3D^. Furthermore, the W285T mutation also increased exchange kinetics in the activation loop (peptide 6: 402-408) and most of the N-terminal lobe, including the αC helix, of Src’s catalytic domain. Thus, β1/β2 resistance cluster mutations appear to promote a more open, regulatory domain-disengaged global state of Src and a dramatically more dynamic N-terminal lobe in Src’s catalytic domain.

The large number of peptic peptides that displayed faster exchange kinetics in W285T Src^3D^ relative to WT Src^3D^ made it difficult to discriminate which differences resulted from localized effects on dynamics versus those that arose from a more open and regulatory domain-disengaged conformational state of Src^3D^. Therefore, we performed a comparative HDX-MS analysis with Src constructs (W285T Src^CD^ and WT Src^CD^) that lack the SH2 and SH3 domain regulatory apparatus. We observed that the W285T mutation led to faster exchange kinetics for a comparable region of Src’s catalytic domain in Src^CD^ to that of Src^3D^ (**Figure 5C**). As in Src^3D^, the peptic peptide that contains the W285T mutation (peptide 3) demonstrated markedly increased exchange kinetics in W285T Src^CD^ relative to WT Src^CD^. In addition to peptic peptide 3, we found that the peptic peptides comprising the remainder of the catalytic domain’s N-terminal lobe (**Figure 5C**) also demonstrated greatly increased exchange kinetics in W285T Src^CD^, suggesting that this mutation directly increases the dynamics of this entire region. Interestingly, a peptic peptide spanning Src’s αC helix (peptide 4: 311-325) demonstrated a large difference in dynamics between W285T Src^CD^ and WT Src^CD^. Given the requirement that Src’s αC helix must adopt an ordered ‘out’ conformation for the catalytic domain to form a high affinity interaction with the SH2-catalytic domain linker and the SH3 domain in the autoinhibited form of Src, the ability of activating mutations in the β1/β2 resistance cluster to directly increase the dynamics of this region provides a plausible mechanism for the reduced intramolecular regulatory domain engagement we observed in our biochemical assays (**Figure 4**). Together, our results support a model where residues in the β1/β2 resistance cluster promote autoinhibitory interactions by restricting the conformational flexibility of the N-terminal lobe, including the αC helix. Activating mutations in the β1/β2 resistance cluster increase the dynamics of the N-terminal lobe, resulting in reduced autoinhibition and a subsequent increase in phosphotransferase activity.

### Mutations in the β1/2 resistance cluster affect the dynamics of Src’s P-loop

Residues in the β1/β2 resistance cluster are part of a structured region that flanks Src’s phosphate-binding loop (P-loop, residues 277-282); a flexible catalytic element that coordinates the γ-phosphate of ATP (**Figure 6A**). A previous study showed that a salt bridge between β1/β2 resistance cluster residues K252 and E260 in the Src Family Kinase (SFK) Lyn (equivalent to K275 and E283 in Src, respectively) appears to affect the dynamics of its P-loop and that mutations that disrupt this electrostatic interaction modulate Lyn’s phosphotransferase activity (Barouch-Bentov et al., 2009). All SFKs contain amino acids that can potentially form a salt bridge at positions equivalent to K275 and E283 in Src (**Figure 6B**) and we observed that, like Src^myr^, a salt bridge-disrupting mutation (E260M) increased the activity of Lyn^myr^ relative to its WT variant in yeast (**Figure 6C, S5A**). Consistent with the potential importance of the salt bridge between K275 and E283 in influencing Src’s phosphotransferase activity, we found that Src constructs that contain a mutation (E283D) that preserves, or potentially enhances, an electrostatic interaction with K275 showed reduced catalytic activity in yeast (**Figure 6C, S5A**) and in *in vitro* activity assays with purified Src constructs (**Figure 6D**) relative to WT. The reduction in phosphotransferase activity of E283D Src^3D^ correlated with increased levels of regulatory domain engagement (**Figure S5B**). Together, previous data and our results suggest that a conserved salt bridge located within the β1/β2 resistance cluster affects the dynamics of the P-loops of SFKs, which influences their phosphotransferase activities.

**Figure 6.**
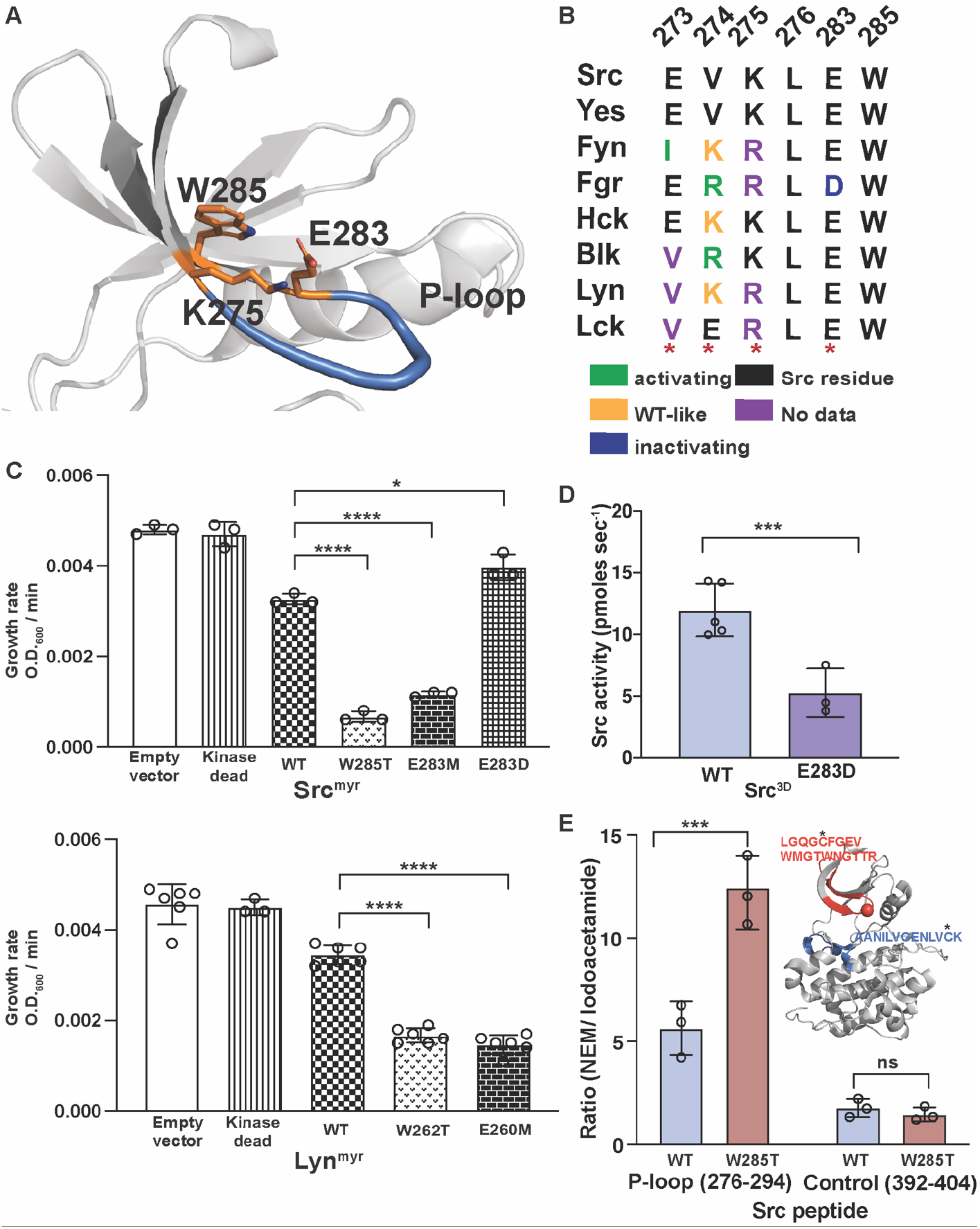
β1/β2 resistance cluster mutations influence the dynamics of Src’s P-loop. (A) Structure of the N-terminal lobe of Src’s catalytic domain (PDB id: 1Y57) showing the flexible region of Src’s P-loop in blue and K275 (β1 strand), E283 (β2 strand), and Trp285 (β2 strand) as orange sticks. (B) Sequence alignment of β1/β2 resistance cluster residues in the SFKs. Classification of activating, WT-like and inactivating mutants derived from (Ahler et.al. 2019). (C) Growth rates of yeast expressing either (*top*) Src^myr^ (Kinase dead, WT, W285T, E283M, or E283D, n = 3) or (*bottom*) Lyn^myr^ (Kinase dead, WT, W262T, or E260M, n = 3-6). (D) Phosphotransferase activity of purified WT and E283D Src^CD^ (n=3-5, \*\*\**P < 0*.*001*). (E) Peak intensity ratios of N-ethylmaleimide (NEM) to iodoacetamide labeling of a Cys-containing tryptic peptide from Src’s P-loop (n=3, \*\*\**P < 0*.*001*) and a Cys-containing control tryptic peptide in Src’s C-terminal lobe (n=3, ns = non-significant, *P > 0*.*05*) from purified WT and W285T Src^CD^. The inset shows the structure of Src’s catalytic domain (PDB id: 1Y57) with the Cys-containing P-loop peptide and its sequence shown in red and the Cys-containing control peptide and its sequence shown in blue. Labeled Cys residues are shown as spheres. See also Figure S5.

We speculated that activating mutations at other positions in the β1/β2 resistance cluster could also influence the dynamics of Src’s P-loop like salt bridge-disrupting mutations. Indeed, our HDX-MS results demonstrated that the two peptic peptides that span the entirely of Src’s P-loop showed greatly increased exchange dynamics in W285T Src^3D^ and Src^CD^ (**Figure 5C, 5D**). However, the large size of the peptic peptides obtained in our HDX-MS workflow made it difficult to determine whether the increase in dynamics we observed was in the P-loop and/or in the structured β-sheet comprising Src’s N-terminal lobe. Therefore, we used a chemical labeling methodology (Potter *et al*., 2020) to probe the dynamics of a cysteine residue (Cys280) located in the center of Src’s P-loop. Specifically, we compared the rates of N-ethylmaleimide (NEM) labeling of Cys280 in WT Src^CD^ and W285T Src^CD^. Consistent with the notion that activating mutations in the β1/β2 resistance cluster modulate the dynamics of Src’s P-loop, Cys280 in W285T Src^CD^ showed a significantly increased rate of NEM labeling relative to WT Src^CD^ (**Figure 6E**). Thus, activating mutations in the β1/β2 resistance cluster not only increase the overall dynamics of the catalytic domain’s N-terminal lobe but also increase the dynamics of the P-loop. We speculate that residues in the β1/β2 resistance cluster act as a network that couples the dynamics of the P-loop to autoinhibitory SH2/SH3 domain interactions that occur on the opposite face of Src’s catalytic domain.

## DISCUSSION

Here, we describe the first saturation mutagenesis study of drug resistance in a multi-domain protein kinase– Src. To do this, we leveraged a yeast-based assay that allowed us to compare how mutations affect Src’s phosphotransferase activity in the absence of inhibitors with its activity in the presence of various ATP-competitive inhibitors. We observed that most inhibitor resistance mutations either conferred a WT-like or activating effect on Src’s phosphotransferase activity in the absence of an inhibitor, which is consistent with the necessity of Src maintaining sufficient catalytic activity in order to obtain drug resistance. We also found that mutations at many residues that the line the ATP-binding pocket of Src’s catalytic domain were capable of conferring resistance to inhibition. Our results with a matched panel of conformation-selective, ATP-competitive inhibitors highlight that many sites within the ATP-binding site where drug resistance occurs are inhibitor independent but that the conformational rearrangements required for certain modes of ATP-competitive inhibition raises opportunities for the emergence of unique mechanisms of resistance. Despite the large number of resistance mutations that line the ATP-binding site of Src, inhibitor contact residues did not represent the most general sites of resistance formation. Instead, residues that participate in the autoinhibition of Src were particularly prone to the development of resistance and mutations that both reduced inhibitor affinity and increased Src’s phosphotransferase activity provided the strongest levels of resistance.

Numerous mutations in a spatially defined cluster of six residues located on the solvent-exposed face of the N-terminal lobe of Src’s catalytic, which we refer to as the β1/β2 resistance cluster, were capable of conferring strong drug resistance. The ability of mutations within the β1/β2 resistance cluster to both generally diminish inhibitor affinity and increase Src’s phosphotransferase activity was surprising given its location on Src’s catalytic domain. The sidechains of all but one (Leu276) of the six residues in β1/β2 resistance cluster are directed away from Src’s ATP-binding site and the top face of Src’s N-terminal lobe has not previously been characterized as a regulatory interface. We found that residues within the β1/β2 resistance cluster appear to be required for limiting the conformational dynamics of Src’s N-terminal lobe and that mutations at these residues release the autoinhibitory interactions required to downregulate Src’s phosphotransferase activity. Thus, residues within the β1/β2 resistance cluster act as a network that couples regulatory domain engagement to the conformational rearrangements in the catalytic domain required for autoinhibition. Mutations that decrease inhibitor residence time by increasing dynamics have been identified as a pathway of resistance in the closely-related tyrosine kinase Abl (Lyczek et al., 2021; Rangwala et al., 2022), and it is likely that mutations in the β1/β2 resistance cluster would have a similar effect on inhibitor residence time for SFKs given their influence on the dynamics of the catalytic domain. Furthermore, our biochemical analyses suggest that the β1/β2 resistance cluster also modulates the dynamics of Src’s P-loop. Residues within the P-loop of Abl are major sites of resistance to inhibitors of the oncogenic fusion protein BCR-Abl in the clinic (**Figure S5C, D**) (Redaelli et al., 2009; Seeliger et al., 2009). However, we observed very few inhibitor resistance mutations in Src’s P-loop because almost all substitutions at these residues were found to be inactivating in our yeast growth assay (**Figure S5E**). Thus, the ability of activating β1/β2 resistance cluster mutations to modulate the conformation of Src’s P-loop while enhancing its phosphotransferase activity may provide a route for acquiring drug resistance that does not require mutation of the P-loop itself. Together, our studies highlight the insights into kinase dynamics and regulation that can be obtained by performing systematic analysis of drug resistance.

## SIGNIFICANCE

Proteins kinases have demonstrated a remarkable ability to acquire mutations that render them less susceptible to inhibition by ATP-competitive inhibitors. Despite the cataloguing of numerous resistance mutations with model studies and in the clinic, a comprehensive understanding of the mechanistic basis of kinase drug resistance is still lacking. To gain a greater understanding of drug resistance in a multi-domain protein kinase, we used a yeast-based screen to determine how almost every mutation in Src’s catalytic domain affects its ability to be inhibited by ATP-competitive inhibitors. Using our yeast-based assay, we compared how mutations affect Src’s phosphotransferase activity in the absence of inhibitors with its activity in the presence of various ATP-competitive inhibitors. As expected, we observed that most inhibitor resistance mutations had a WT-like or activating effect on Src’s phosphotransferase activity in the absence of inhibitors. We also found that a number of mutations at positions that the line the ATP-binding site of Src provided general resistance to ATP-competitive inhibitors. However, a subset of resistance mutations appear to be specific to the structural rearrangements required for certain modes of ATP-competitive inhibition. While inhibitor contact residues represented sites of resistance, we found that residues that participate in the autoinhibition of Src are particularly prone to the development of resistance. Biochemical analysis of a resistance-prone cluster of residues revealed that the top face of Src catalytic domain’s N-terminal lobe, unexpectedly, contributes to the autoinhibition of Src and that mutations in this region led to resistance by lowering inhibitor affinity and promoting kinase hyperactivation. Together, our studies demonstrate how comprehensive profiling of drug resistance can be used to understand potential resistance pathways and uncover new mechanisms of kinase regulation.

## ACKNOWLEDGEMENTS

This work was supported by the National Institute of General Medical Sciences (R01GM109110 to D.M.F. and R01GM086858 to D.J.M.) and National Human Genome Research Institute (RM1HG010461 to D.M.F).

## AUTHOR CONTRIBUTIONS

S.C. and E.A. conducted experiments and performed data analysis. S.C. and M.G. designed and performed HDX-MS experiments. J.J.S, L.F., Z.E.P., K.A.S., and J.J.S. generated reagents and performed experiments. S.C., E.A., D.M.F., and D.J.M. conceived of the project, designed experiments, and wrote the manuscript.

## KEY RESOURCES TABLE

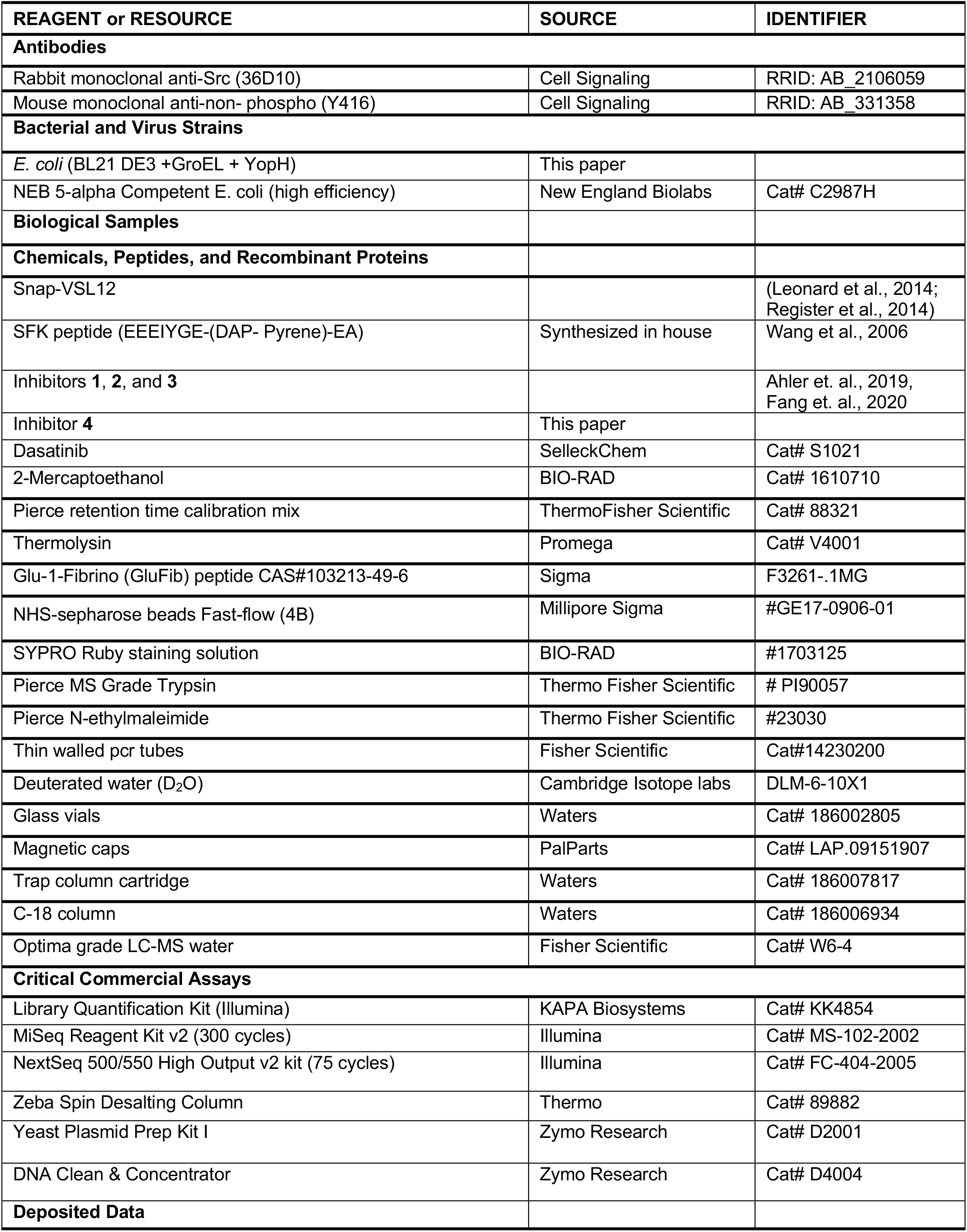
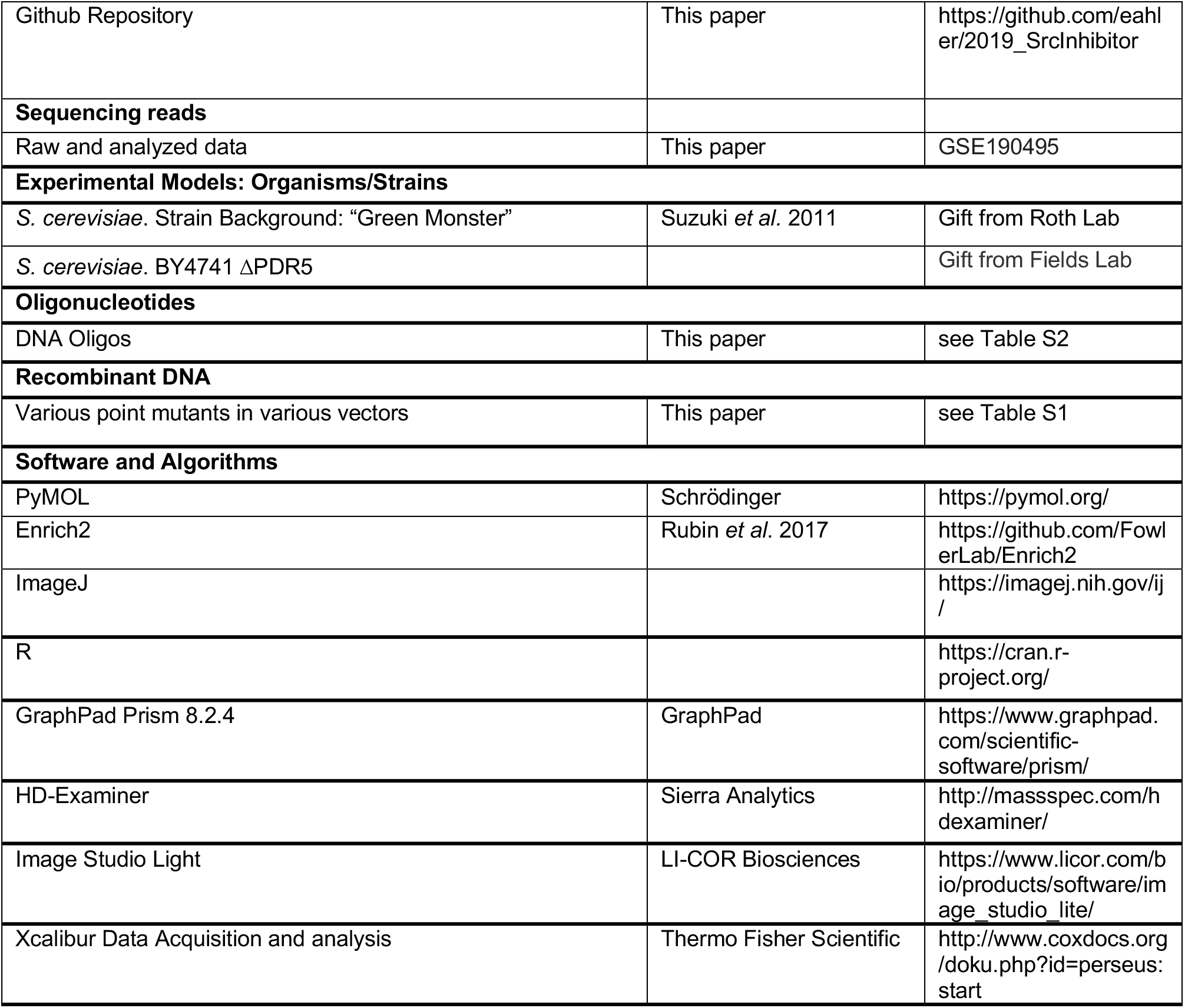

## STAR METHODS

### RESOURCE AVAILABILITY

#### Lead Contact

Further information and requests for resources and reagents should be directed to and will be fulfilled by the Lead Contact, Dustin J. Maly.

#### Materials availability

Plasmids generated in this study are available upon request from the lead contact.

#### Data and code availability

DMS data have been deposited at GEO and will be made publicly available as of the date of publication. Accession numbers and links to all python and R code have been listed under the section Additional Resources and Key Resource Table.

### EXPERIMENTAL MODEL AND SUBJECT DETAILS

#### *S. cerevisiae* genetics and cell culture

BY4741 Green Monster (Suzuki et al., 2011; a generous gift from Dr. Fritz Roth, inhibitors **1, 2** and **4)** or BY4741 (ΔPDR5, MATa His3D1 Leu2D0 Met15D0 Ura3D0, dasatinib) were used to perform yeast experiments. All Src constructs were cloned into the p415GAL1 plasmid. To select successful transformants, yeast was transformed using standard LiAc protocols (Gietz and Schiestl, 2007) and plated on C-Leu media. All growth experiments were performed in C-Leu media to maintain plasmid.

#### Bacterial cell culture

*In vitro* experiments were performed with recombinant protein purified from thermo-competent *E*.*Coli* (BL21 DE3) cells expressing YopH and GroEL (prepared in house).

## METHOD DETAILS

Restriction enzymes were purchased from New England Biolabs and all chemicals purchased from Sigma unless otherwise specified.

### Cloning

QuickChange Site Directed Mutagenesis (Agilent) or IVA cloning (Garcia-Nafria et al., 2016) was used to generate all mutants discussed in the paper following standard protocols. Mutations were verified by Sanger sequencing of the entire open reading frame. Gibson assembly or directional cloning following standard protocols was performed to achieve all subcloning and all constructs were validated by Sanger sequencing.

### Western blotting

Src antibody (36D10; CST #2109) and Non-phospho-Src (Tyr416) (7G9; CST #2109) were purchased from Cell Signaling Technology (CST). Anti-rabbit secondary antibody was purchased from Li-Cor.

### Protein purification and expression

For *in vitro* biochemical assays, Src^FL^ (WT/E283M/V326K/W285T, residues 2-536), Src^3D^ (WT/W285T/E283M/E283D, residues 87-536), Src^CD^ (WT/W285T/E283M/E283D, residues 261-536) and autoinhibited Src^EEI^ (WT/W285T, residues 87-536; mutations Q531E, P532E and G533I) were cloned into the bacterial expression plasmid pMCSG7 as N-terminal His_6_-SUMO tagged constructs. Src constructs were co-transformed into *E*.*coli* expressing YopH/GroEL and plated on triple selective plates (Ampicillin/Chloramphenicol/Streptomycin). A single colony was picked and grown in an overnight culture of 15 mL of Terrific broth containing all three antibiotics. A 1 L culture was then inoculated with the starter culture, grown to an O.D._600_ of 1.1, the temperature was then dropped to 18*°*C and protein expression was induced with 0.4 mM IPTG overnight. Ni-NTA was used to purify His_6_-SUMO-Src after lysing cells in lysis buffer (50 mM HEPES, pH 8.0, 300 mM NaCl, 1 mM PMSF, 0.1% Triton-X, 20 mM imidazole) and eluted using purification buffer (50 mM HEPES, pH 8.0, 300 mM NaCl, 1 mM PMSF, 0.1% Triton-X, 10% glycerol, 0.2% BME, 150 mM imidazole). A 2 h dialysis in dialysis buffer (50 mM HEPES, pH 8.0, 150 mM NaCl, 1 mM DTT, 10% glycerol) was performed at 4*°*C prior to adding the SUMO protease His_6_-ULP1 (1:25 protease:eluted protein, wt/wt). The Src-protease mixture was then transferred to a fresh dialysis buffer and was allowed to cleave overnight at 4*°*C. Following cleavage, a second Ni-NTA purification was carried out to remove any non-cleaved Src and His_6_-ULP1. Finally, an anion exchange column (Pierce, 90011) was used to remove YopH and GroEL to yield Src at > 95% purity. Relates to Figure 3F, S3F, 4B, 4D-I, S4F-J, 5C-D, S5, 6D-E, S5B

### Src phosphotransferase activity

Purified recombinant Src variants were used to measure Src phosphotransferase activity using a self-reporting fluorescent SFK peptide (EEEIYGE-(DAP-Pyrene)-EA) (Wang et al., 2006) in an *in vitro* kinase assay. Briefly, 20 μL of purified Src^FL^ (6.5 nM), Src^3D^ (5 nM), Src^CD^ (5.5 nM) or autoinhibited Src^EEI^ (8 nM) constructs of each variant (WT, V326K, W285T, E238M or E283D) were diluted in kinase reaction buffer (76 mM HEPES, pH 7.5, 5 mM MgCl_2_, 150 mM NaCl, 3.8 mM EGTA, 0.2 mg/mL BSA, 150 μM Sodium orthovanadate (Na_3_VO_4_)) and incubated with 5 μL 1 mM ATP at room temperature for 30 min. Next, 5 μL of 40 μM SFK peptide was added to each well and raw fluorescence units were measured immediately on an Envision fluorometer (Perkin Elmer) with an excitation wavelength of 344 nm and an emission wavelength of 405 nm in real time for the first 15 min at 15 s intervals. Calculation of kinase activity in terms of pmole s^-1^ of phosphorylated substrate per nM of enzyme is discussed below under “Calculation of Src activity.” Relates to Figure S3F, 4D-F, 6D.

### Src *K*_*M*_ [ATP] determination

*K*_*M*_ [ATP] of purified Src variants were measured using the same assay described above. Briefly, 2-fold (8-data points) serial dilution of ATP starting at 1 mM, was incubated with 5 nM of Src (WT or mutant) in kinase reaction buffer (76 mM HEPES, pH 7.5, 5 mM MgCl_2_, 150 mM NaCl, 3.8 mM EGTA, 0.2 mg/mL BSA, 150 μM Na_3_VO_4_) and 20 μM of SFK peptide. Raw fluorescence units were measured immediately on an Envision fluorometer (Perkin Elmer) with an excitation wavelength of 344 nm and an emission wavelength of 405 nm in real time for 90 mins at 15 min intervals. Calculation of kinase Km [ATP] is discussed below under “Calculation of IC_50_, *K*_*M*_, *K*_*I*_.” Relates to Figure S3F, S4G.

### Inhibitor IC_50_ and K_i_ determination

For all IC_50_ determination experiments, first a kinase titration was performed as described above prior to inhibitor titration to ensure linearity of kinase concentration in the assay. Inhibitors (initial concentration = 30 μM, 3-fold serial dilutions, 10 data points in triplicate) were assayed against Src^FL^ ([WT] = 4 nM ; [W285T] = 5 nM; [V326K] = 5 nM; [E283M] = 4.5 nM), and Src^CD^ ([WT] = 3.5 nM, [W285T] = 5 nM;

[E283M] = 4.5 nM) in assay buffer (76 mM HEPES, pH 7.5, 5 mM MgCl_2_, 150 mM NaCl, 3.8 mM EGTA, 0.2 mg/mL BSA, 150 μM Na_3_VO_4_). Briefly, kinase was pre-incubated with 1 mM ATP and inhibitors for 30 min in a 384-black assay plate (Corning, #3573). 20 μM of SFK peptide was then added to plate and incubated for 2 h. Raw fluorescence units were measured on Envision (Perkin Elmer) with excitation wavelength of 344 nm and emission wavelength of 405 nm. Data was analyzed using GraphPad Prism 8.4.2 software, IC_50_ determination and *K*_*I*_ calculation are discussed below under “Calculation of IC_50_, *K*_*M*_, *K*_*I*_.” Relates to Figure 3F, 4B.

### SFK yeast growth assay

SFK yeast growth assay was performed as described in (Ahler et al., 2019). Briefly, codon-optimized full-length human Src or Lyn (WT or indicated mutants) were transformed into the *S. cerevisiae* BY4741 Green Monster or BY4741 (ΔPDR5) strain using standard LiAc transformation protocols (Gietz and Schiestl, 2007) and plated on C-Leu plates to select for successful transformants. Two or three independent colonies for each strain were collected and treated as biological replicates. Single colonies for each strain were grown overnight in 5 mL of 3% raffinose C-Leu to saturation. The following day cultures were back diluted to O.D._600_ = 0.5 in C-Leu 3% raffinose and grown to at least O.D._600_ = 1.0 before subsequent dilution. To induce expression, cultures were diluted to O.D._600_ = 0.01 into C-Leu 2% galactose, then 150μL of each culture was plated and grown in a BioTek Synergy plate reader under constant shaking at 30*°*C. O.D_600_ was measured every 30 min over a 36 or 48 h period.

For SFK yeast growth assays with inhibitors, the same procedures were followed as above except the indicated inhibitor concentration (or DMSO vehicle control) was added at the time of induction (final DMSO = 1%). To calculate the growth rate for an individual variant, background corrected O.D._600_ values within the range 0.04-0.32 from the yeast growth assay, corresponding to 2-5 doublings, were used. During this phase of growth, the effects of culture density and detector signal sensitivity on yeast growth rate were negligible, ensuring a linear growth pattern. The background-corrected O.D._600_ values were natural log-transformed and the slope of the line for time vs. ln(O.D._600_) were calculated (reported as the growth rate). Relates to Figure 1, S1, 2, S2, 3B-E, S3A-E, S4B-E, 6C, S5A.

### Generation of Src variant library for inhibitor screening

The Src variant library was generated as previously described in (Ahler et al., 2019). Drug resistant screens for dasatinib were performed in the BY4741 strain and for inhibitors **1, 2**, and **4** in BY4741 Green Monsters. Selection in presence of inhibitors was initiated by inoculation at a final of O.D._600_ 0.01 into 200 μL C-Leu 2% galactose media (dasatinib (25 or 100 μM), inhibitor **1** (8 μM), inhibitor **2** (2 μM), inhibitor **4** (0.8 μM), or DMSO were added during this step). Growth was monitored throughout the selection by measuring O.D_600_. Selection O.D._600_s are as follows; for 25 μM dasatinib: 0.22, 2.7, 11.7 (replicate #1) and 0.18, 2.1, 12.04 (replicate #2); 100 μM dasatinib: 0.26, 3.1, 11.6 (replicate #1) and 0.19, 2.2, 9.8 (replicate #2); for 8 μM of inhibitor **1**: 0.22, 0.65, 2.5 (replicate #1) and 0.39, 1.5, 5.2 (replicate #2); for 2 μM of inhibitor **2**: 0.18, 0.62, 4 (replicate #1) and 0.19, 0.83, 3.6 (replicate #2); for 0.8 μM of inhibitor **4**: 0.15, 1.2, 6.9 (replicate #1) and 0.32, 1.1, 3.6 (replicate #2). Yeast samples from all O.D._600_ listed were harvested and pelleted at 3000xg for 5 min and pellets were frozen at -80 °C. Next, plasmids were extracted from frozen pellets using Yeast Plasmid Prep I (Zymogen) according to manufacturer’s protocol and resuspended in 15 mL H_2_O. To append Illumina cluster generators and append indices, 25 μL 2x KAPA2G Robust HotStart ReadyMix (KAPA Biosystems), 2 μL 10 mM SC10-SC12 (indexing primers), 2 μL 10 mM SC13-SC17 (indexing primers), 13.5 μL H_2_O, and 7.5 μL extracted plasmid. Thermocycler conditions used were: Initial denaturation at 95 °C for 3 min, followed by 17 cycles of 95 °C for 15 s, 60 °C for 15 s, and 72 °C for 15 s. Each amplification was cleaned, quantified using KAPA Library Quantification Kit, then sequenced on NextSeq 500/550 High Output v2 kit (75 cycles) with standard Illumina sequencing primers and the custom primers SC18 and SC19.

### Src variant library analysis

Src variant library analysis was performed as previously described in (Ahler et al., 2019)

### Identifying resistance mutations

The Src variant library was treated with dasatinib or various conformation-selective inhibitors and resistance mutations were identified based on variant activity score. Upon inhibitor treatment, yeast harboring drug sensitive Src variants have their growth rescued, while those expressing drug resistant Src variants continue to grow poorly. Time points were sampled throughout growth, plasmids extracted, barcodes amplified and deeply sequenced on an Illumina NextSeq run as described above. We calculated the mean and standard deviation of activity scores for all synonymous variants (for dasatinib STD= 0.35, for inhibitors **1, 2** and **4** STD = 0.33, 0.31 and 0.33, respectively) identified in our dataset. All variants were classified as ‘resistant’ if their activity score were 2x of this value (= 0.7 for dasatinib, =0.66 for **1**, =0.62 for **2**, and =0.66 for **4**).

### SH3 domain pulldown assays

SH3 pulldown experiment was performed as described previously in (Ahler et al., 2019). Briefly, 20 μL of a 50% slurry of SNAP-capture pulldown resin (prepared using NHS-sepharose beads Fast-flow (4B)) was placed in a micro-centrifuge tube. The resin was washed (3x, 10 bead volumes) with pulldown buffer (20 mM Tris-HCl, pH 7.5, 100 mM NaCl, 1 mM DTT and 0.2 mg/mL BSA). 8 μM of SNAP tag–polyproline peptide fusion (VSLARRPLPPLP) was loaded onto the resin at a final volume of 50 μL per 10 μL of bead in pulldown buffer (Leonard et al., 2014). The resin was equilibrated at room temperature for 1 h and then washed (3x, 10 bead volumes) prior to performing pulldown assays.

Next, SH3 pulldown assays performed with recombinant 100 nM of Src^3D^ (WT or W285T) or Src^EEI^ (WT or W285T) in 50 μL of pulldown buffer, incubated with 5 μL of the immobilized SH3 domain ligand. The resin-Src mixture was equilibrated room temperature for 1 h on a rotator. Resin was spun down using a mini centrifuge, the supernatant was aspirated, and the resin was then washed three times before eluting the retained kinase with 50 μL of 1x SDS loading buffer. The beads were boiled at 95 *°*C for 10 min. All samples were separated via SDS–PAGE and visualized by western blotting with Src antibody (Cell Signaling, #2109 for WT and #2102 for Src^EEI^) on Li-Cor Odyssey. The scanned blots were quantified with ImageStudio Lite software and the signal corresponding to input protein (“I”) was scaled to the original loaded kinase amount and signal corresponding to eluted (“E”) was measured to determine kinase retained on the resin (% retained Src) based on the loaded and eluted fraction based on curve fitting of immunoblot signal intensity to a Src titration. Relates to Figure 4G, S4H.

### Limited proteolysis of Src with thermolysin

Methods for limited proteolysis of Src was modified from (Agius et al., 2019). Briefly, Src^3D^ (WT, W285T, E283M, or E283D) or autoinhibited Src^EEI^ (WT, E283M, or W285T) was diluted to 1 μM in proteolysis buffer (50 mM Tris-HCl pH 8.0, 100 mM NaCl, 0.5 mM CaCl_2_). Proteolysis was initiated by adding a 3.8 mM Thermolysin (Promega, #V4001) stock solution to the kinase (final concentration of thermolysin = 60 nM). 20 µL of this mixture was then added to 10 µL of 50 mM EDTA in 1x loading buffer to terminate proteolysis at various time points (0, 2, 4, 8, 16, 32, 64, 128, 256 min). The quenched samples were analyzed by SDS-PAGE (12% Bis-Tris gel in SDS running buffer) and stained with SYPRO Ruby (ThermoFisher Scientific: #S12000) according to the manufacturer’s protocol. Band intensities were analyzed by ImageStudioLite imaging software. Percent protein remaining was computed as relative Src band intensity at 0 min and was plotted against time on GraphPad Prism 8.4.2. The curve was fit to an exponential decay equation using GraphPad Prism 8.4.2 software to obtain the half-life of each Src variant. Relates to Figure 4H-I, S4I, S4J, and S5B.

### HDX-MS of Src

HDX-MS of Src was performed as describe in (Potter et al., 2020). Briefly, purified Src^3D^ (WT or W285T) or Src^CD^ (WT or W285T) was diluted to 0.2 mg/mL in protein dilution buffer (50 mM HEPES, pH 7.8, 150 mM NaCl, 1 mM DTT, 5% glycerol). 10 μL of this dilution was then added to 90 μL of buffered D_2_O (prepared 5 mL with 4.5 mL of D_2_O and 0.5 mL of 10x protein dilution buffer and 0.2 μg/mL of peptide standard Glu-1-Fibrino peptide (CAS: 103213-49-6, Sigma)) to initiate deuteration at 22 °C. Deuterium exchange was quenched after 3 s, 1 min, 30 min, and 20 h by adding the reaction to 100 μL of ice-cold quench buffer (0.2% formic acid, 8M Urea, 0.1% trifluoroacetic acid, allowing final pH to drop to 2.5) in order to lock deuterium in place and unfold the protein. All time points were collected in triplicate (except the 20 h samples of Src^CD^ which was collected in duplicate). Samples were immediately frozen in a dry ice/ethanol bath and stored at –80°C until LC-MS analysis. Undeuterated samples were prepared the same way except using buffered H_2_O (Optima grade LC-MS water, Fisher Scientific, product #W6-4) instead of D_2_O. Frozen samples were thawed on a 5 °C block for 4 min prior to injection onto a loading loop. The loaded sample was passed over a custom packed pepsin column (Porcine pepsin immobilized on POROS 20-AL resin; 2.1 × 50 mm column) (Wang et al., 2002) kept at 12 °C with a flow of 0.1% trifluoroacetic acid (TFA) and 2% acetonitrile (ACN) at 200 µL/min. Digested peptic fragments were trapped onto a Waters XSelect CSH C18 XP VanGuard Cartridge (2.1 × 5 mm, 2.5 µm). After 5 minutes of loading, digestion, and trapping, peptides were resolved on an analytical column (Waters C18 BEH 1 × 100 mm, 1.7µm, 130Å) using a gradient of 3% to 40% solvent B for 9 minutes (A: 0.1 % FA, 0.025 % TFA, 2 % ACN; B) 0.1% FA in ACN). The LC system was coupled to a Thermo Orbitrap performing full scans over the *m/z* range of 300 to 1500 at a resolution of 30,000. The MS source conditions were set to minimize loss (Walters et al., 2012). Undeuterated samples were run prior to and at the end of all the LC-MS queues.

During the analytical separation step, a series of 250 µL injections were used to clean the pepsin column: 1) 0.1% Fos-12 with 0.1% TFA; 2) 2 M GndHCl in 0.1% TFA; 3) 10% acetic acid, 10% acetonitrile, 5% isopropanol (Majumdar et al., 2012) (Hamuro and Coales, 2018). After each gradient the trapping column was washed with a series of 250 µL injections: 1) 10% FA; 2) 30% trifluoroethanol; 3) 80% methanol; 4) 66% isopropanol, 34% ACN; 5) 80% ACN. During the trap washes the analytical column was cleaned with three rapid gradients (Fang et al., 2011).

Peptic peptides were identified from data-dependent acquisition (DDA) experiments on undeuterated samples by exact mass and tandem mass spectrometry (MS/MS) spectra using Protein Prospector (Baker and Chalkley, 2014) filtering with a score cutoff of 15. Mass shifts were determined using HD-Examiner v2 (Sierra Analytics). The Glu-1-Fibrino internal standard peptide was checked in all samples to verify that back-exchange levels were consistent in all experiments (Zhang et al., 2012). Peptic petides with significant differences in exchange were assessed using the hybrid significance threshold at 99% CI (Table S5) as well as consistency among all observed overlapping peptides (Hageman, et al., 2019). Based on this the difference cut-off was set at 0.557 Da in case of Src^3D^ and 0.435 Da in case of Src^CD^ and all peptides that pass this cut-off has been mapped in red onto the crystal structure of Src^3D^. Relates to Figure 5C and Table S5.

List of identified peptides as well as details of HDX exchange has been submitted as a supplemental table 5.

### Maleimide labeling and mass spectrometry

Maleimide labeling and subsequent mass spectrometry analysis of cysteine labeled peptides was performed as described in (Ahler et al., 2019). Briefly, purified 1 μM Src^CD^ (WT or W285T) was diluted in mass spectrometry buffer (50 mM HEPES, pH 7.6, 150 mM NaCl, 5% glycerol and 0.02% (wt/vol) n-Dodecyl β-D-maltoside (DDM)) (Kahsai et al., 2014) and treated with 100 μM N-ethyl maleimide (NEM) in a LoBind 1.5 mL Eppendorf tube at 25 °C for 30 min. The NEM labeling reaction was quenched with 20 mM of DTT in ammonium bicarbonate (NH_4_HCO_3_) solution. Protein was then precipitated using 0.02% deoxycholate and 10% trichloroacetic acid on ice for 10 min. The precipitated protein was pelleted by spinning at 10,000x rpm for 15 min. The pellet was dried with 10 μL acetone and resuspended in peptide solubilization buffer (8 M urea, 200 mM Tris-HCl, pH 8.0, 2.4 mM iodoacetamide (IA), 0.001% DDM) by vortexing briefly. The mixture was incubated in the dark for 30 min. Trypsin digestion solution (0.5 mg/mL of trypsin in 1 mM CaCl_2_, 200 mM Tris-HCl, pH 8.0) was added and the protein was digested overnight at 37°C. Peptide was desalted using C-18 ZipTips (Milipore) and each sample was run on the Finnigan LTQ Ion trap. [M+3H]+3 peptide masses for both NEM and iodoacetamide modified, cysteine-containing peptide was analyzed using Xcalibur MaxQuant software. For each tryptic peptide (LGQGCFGEVW(<u>T</u>)MGTWNGTTR (P-Loop) and AANILVGENLVCK (control)), the ionic intensity for both the NEM-labeled and IA-labeled species were obtained, and a ratio was calculated. The experiment was repeated in triplicate. Relates to Figures 6E.

### Synthesis of inhibitors

General Synthetic Procedures: All chemicals purchased from commercial suppliers were used without further purification unless otherwise stated. Reactions were monitored with thin-layer chromatography (TLC) using silica gel 60 F254 coated glass plates (EM Sciences). Compound purification was performed with an IntelliFlash 280 automated flash chromatography system using pre-packed Varian SuperFlash silica gel columns (Hexane/EtOAc or CH2Cl2/MeOH gradient solvent). A Varian Dynamax Microsorb 100-5 C18 column (250 mm x 21.4 mm), eluting with H_2_O/CH_3_CN or H_2_O/MeOH gradient solvent (+0.05% TFA), was used for preparatory HPLC purification. The purity of all final compounds was determined by analytical HPLC with an Agilent ZORBAX SB-C18 (2.1 mm x 150 mm) or Varian Microsorb-MV 100-5 C18 column (4.6 mm x 150 mm), eluting with either H_2_O/CH_3_CN or H_2_O/MeOH gradient solvent (+0.05% TFA). Elution was monitored by a UV detector at 220 nm and 254 nm, with all final compounds displaying > 95% purity. Nuclear Magnetic Resonance (NMR) spectra were recorded on Bruker 300 or 500 MHz NMR spectrometers at ambient temperature.

Inhibitors **1, 2**, and **3** was synthesized as previously described in (Ahler et al., 2019; Fang et al., 2020).

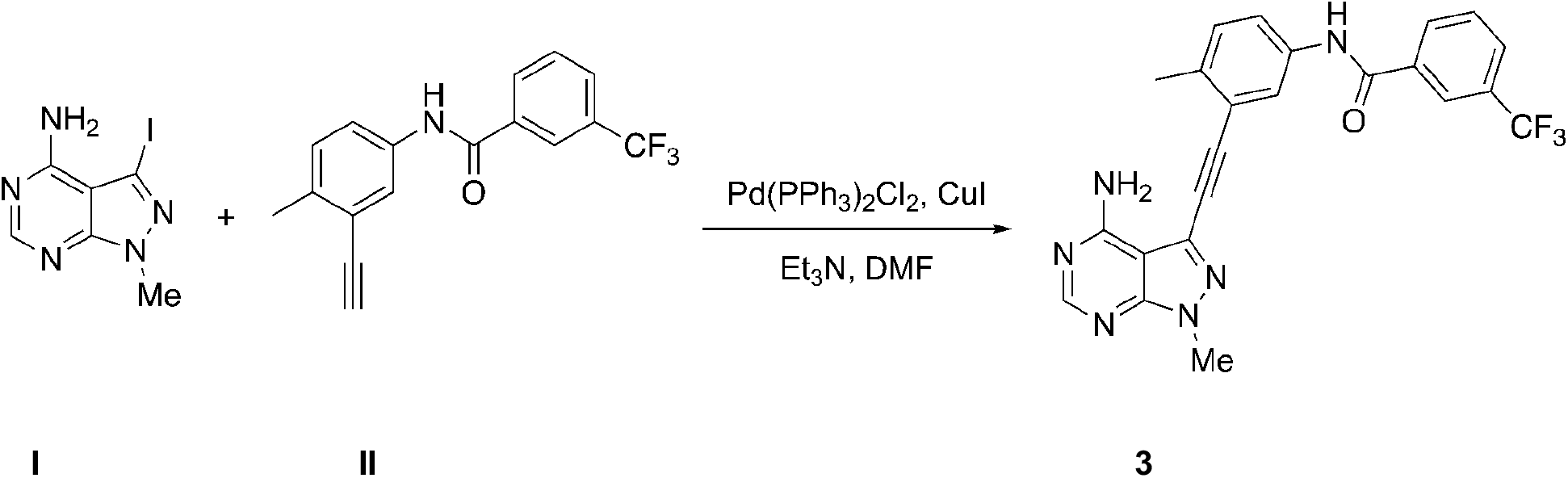

3-Iodo-1-methyl-1H-pyrazolo[3,4-d]pyrimidin-4-amine (**I**, 73 mg, 0.27 mmol, 1.0 equiv.) was dissolved in anhydrous DMF (3.5 mL) under nitrogen. Triethylamine (110 mg, 150 μL, 1.1 mmol, 4.0 equiv.), N-(3-ethynyl-4-methylphenyl)-3-(trifluoromethyl)-benzamide (**II**, 120 mg, 0.40 mmol, 1.5 equiv.), bis(triphenylphosphine) palladium(II) dichloride (9.3 mg, 0.013 mmol, 0.05 equiv.), and copper (I) iodide (5.1 mg, 0.027 mmol, 0.10 equiv.) were added to the above solution sequentially. The reaction was heated at 50 °C for overnight under nitrogen and then quenched with a saturated NH_4_Cl aqueous solution (5 mL). The resulting mixture was diluted with ethyl acetate (40 mL) and the organic phase was washed with a saturated NaHCO_3_ aqueous solution (10 mL), brine (10 mL), and then dried over anhydrous Na_2_SO_4_. Purification by flash chromatography on silica gel using a gradient of EtOAc/Hexane (0:100 to 100:0) afforded inhibitor **3**, N-(3-((4-amino-1-methyl-1H-pyrazolo[3,4-d]pyrimidin-3-yl)ethynyl)-4-methylphenyl) 3(trifluoromethyl)benzamide as a pale brown solid (70 mg, 58%).

^1^H-NMR (300 MHz, DMSO) δ= 10.57 (s, 1H), 8.43 – 8.27 (m, 3H), 8.09 (s, 1H), 8.01 (d, *J* = 7.6 Hz, 1H), 7.92 – 7.74 (m, 2H), 7.39 (d, *J* = 8.5 Hz, 1H), 3.98 (s, 3H), 3.39 (s, 3H), 2.53 (s, 2H); MS (ESI, m/z) calculated for C_23_H_17_F_3_N_6_O 450.1, [M+H]^+^ found 451.1. HPLC purity>99%. Relates to Figure 3-4, S3, S4.

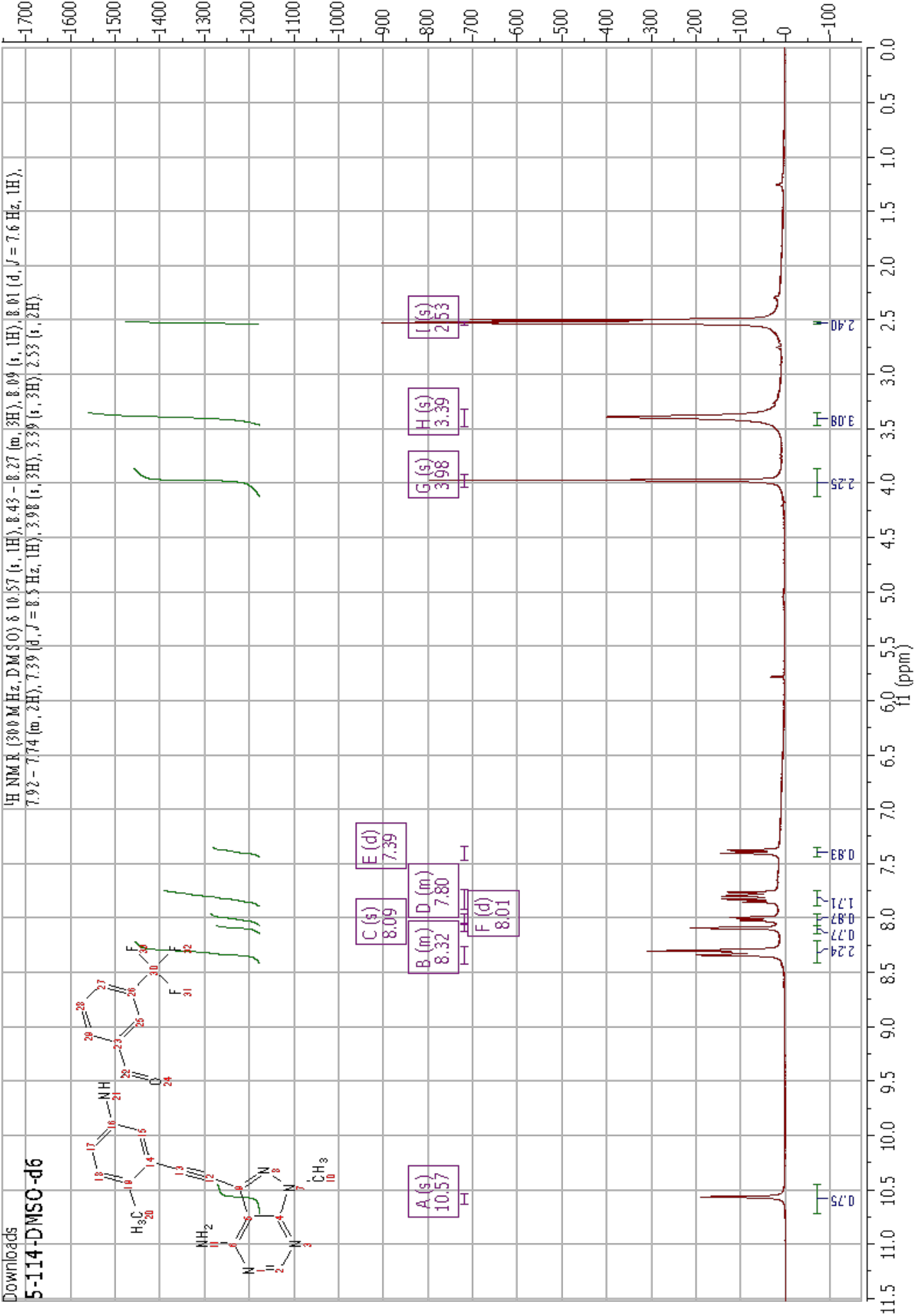

### QUANTIFICATION AND STATISTICAL ANALYSIS

For all statistical tests (unless otherwise noted), a two-tailed Student’s t test was used to compare means between two samples. A one-way ANOVA with post hoc Tukey’s HSD test was used to compare means between more than two samples. Statistical tests were performed in R or GraphPad Prism 8.4.2. SEM for n = 3-6 has been reported for all *in vitro* biochemical assays. Significance is denoted as * = p < 0.05, *** = p < 0.001, **** = p < 0.0001.

### Calculation of *K*_*I*_, IC_50_ and *K*_*M*_

IC_50_ values were calculated in GraphPad Prism using the ‘‘One-Site Fit log IC50.’’ *K*_*M*_ [ATP] values were determined using GraphPad Prism using ‘‘Plot Michaelis-Menten’’ option. *K*_*I*_ values for all Src constructs were calculated using the Cheng-Prusoff equation at 1 mM ATP and calculated *K*_*M*_ [ATP]. Relates to Figures 3F, S3F, 4B, S4G.

### Classification of Src variants

Classification of Src variant library from activity scores as activating, WT-like and inactivating was performed as previously described in (Ahler et al., 2019). Relates to 4C, S2A, S3C, S5C, S5E.

### Calculation of Src phosphotransferase activity

Slopes were calculated from the linear portion of *in vitro* kinase enzyme assay to first obtain raw fluorescence count per sec. This was then divided by fluorescence change per picomoles of phosphorylated substrate (obtained from the slope of a standard curve of raw fluorescence versus phosphorylated substrate) to calculate phosphorylated substrate produced in pmoles sec^-1^ reported per nM of the Src variants tested. Relates to Figures S3F, 4D-F.

## ADDITIONAL RESOURCES

### Data and software availability

Sequencing reads were deposited in NCBI’s Gene Expression Omnibus (GEO) and are accessible through accession number GSE190495. Raw Data for the Src DMS and code to reproduce figures are located in our Github repository at https://github.com/eahler/2019_SrcInhibitor.

Any additional information required to reanalyze the data reported in this manuscript is available upon request from the lead contact.

