## Supplemental Figure and Table for "Profiling of the drug resistance of thousands of Src tyrosine kinase mutants uncovers a regulatory network that couples autoinhibition to catalytic domain dynamics": Supplemental Figures_20230319.pdf

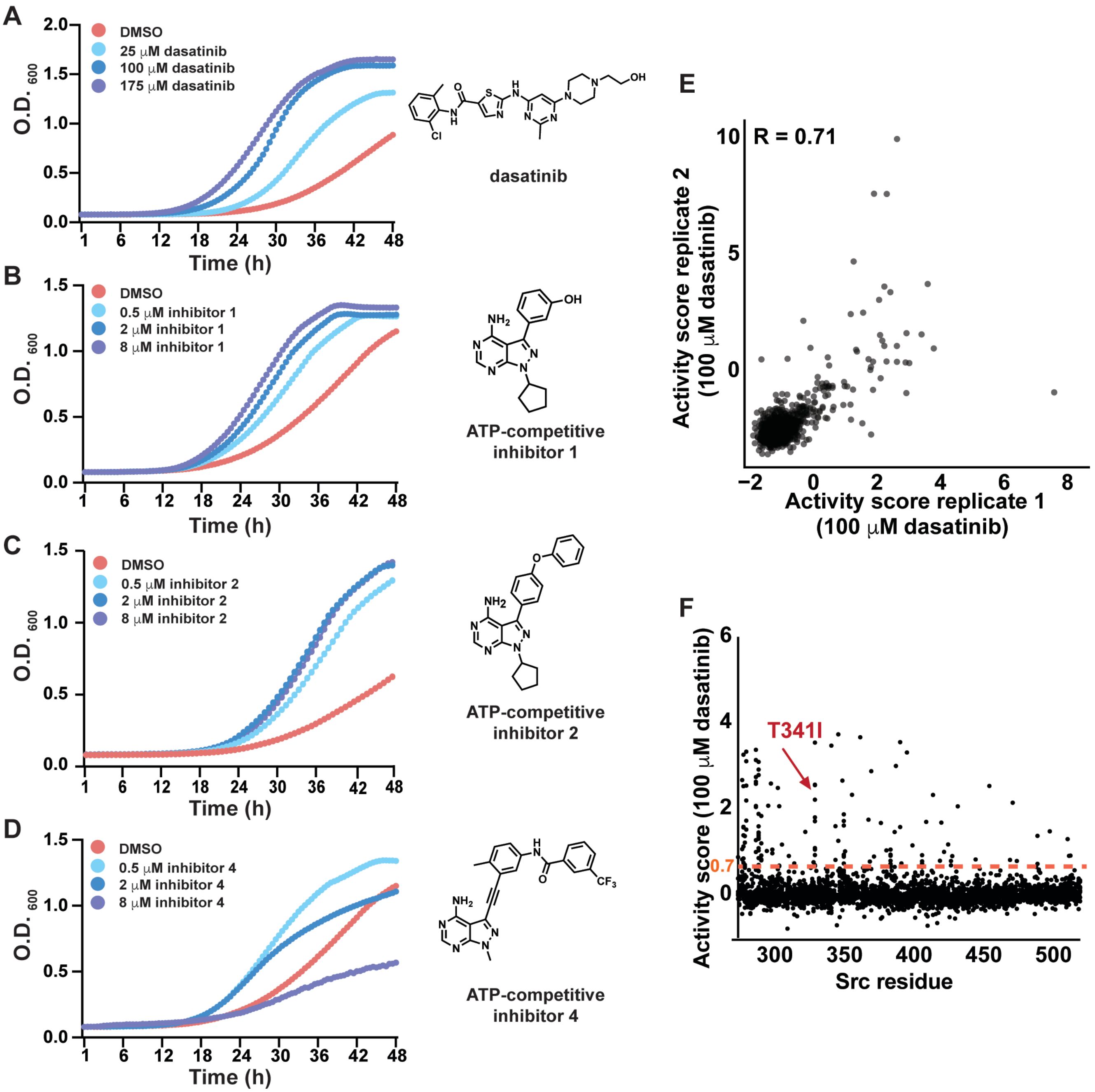


**Figure S1. A yeast-based assay for the parallel analysis of drug resistance in Src tyrosine kinase.** (A) Individually assessed growth curves for yeast expressing wild-type (WT) Src$^{myr}$ in the presence or absence of different concentrations of dasatinib (*left*, n=3). Structure of dasatinib (*right*). DMSO data is derived from Ahler et. al. 2019. (B) Individually assessed growth curves for yeast expressing WT Src$^{myr}$ in the presence or absence of different concentrations of ATP-competitive inhibitor **1** (*left*, n=3). Structure of inhibitor **1** (*right*). (C) Individually assessed growth curves for yeast expressing WT Src$^{myr}$ in the presence or absence of different concentrations of ATP-competitive inhibitor **2** (*left*, n=3). Structure of inhibitor **2** (*right*). (D) Individually assessed growth curves for yeast expressing WT Src$^{myr}$ in the presence or absence of different concentrations of ATP-competitive inhibitor **4** (*left*, n=3). Structure of inhibitor **4** (*right*). (E) Scatterplot showing activity score correlations (Pearson's R = 0.71) between two independent transformations of the Src$^{myr}$ variant library in yeast treated with 100 μM dasatinib. (F) Scatterplot showing activity scores (each black dot represents the average of two replicates) for every single amino acid substitution at every position in Src's catalytic domain for yeast transformed with the Src$^{myr}$ library and treated with 100 μM dasatinib.

| Construct | Expressed in | Residue numbers | Vectors used in | Notes | Citation |
|---|---|---|---|---|---|
| $Src^{myr}$ WT, W285T, E283M, E283D | *S. cerevisiae* | 1-536 | p415 GAL1 | | This study |
| $Src^{myr}$ K298M (kinase dead) | *S. cerevisiae* | / | p415 GAL1 | Catalytically-inactive mutant | Florio *et al.*, 1994 |
| $Src^{FL}$ WT | *E. coli* | 2-536 | pMCSG7 | N-terminal His6-SUMO tag gets cleaved | Ahler *et. al.*, 2019 |
| $Src^{FL}$ V326K | *E. coli* | 2-536 | pMCSG7 | Drug resistant mutant | This study |
| $Src^{FL}$ E283M, E283D, W285T | *E. coli* | / | p415 GAL1, pMCSG7 | β1/ β 2 resistance cluster mutants | This study |
| $Src^{EEI}$ WT | *E. coli* | 87-536 | pMCSG7 Q531E, P532E, G533I | Autoinhibited Src | Chakraborty *et. al.*, 2019 |
| $Src^{EEI}$ E283M, W285T | *E. coli* | 87-536 | pMCSG7 Q531E, P532E, G533I | β1/ β 2 resistance cluster mutants (autoinhibited) | This study |
| $Src^{3D}$ E283M, E283D, W285T | *E. coli* | 87-536 | pMCSG, p415 GAL1 | N-terminal His6-SUMO tag gets cleaved | This study |
| $Src^{CD}$ WT | *E. coli* | 261-536 | pET28a, pMCSG7 | N-terminal His6-SUMO tag gets cleaved | This study |
| $Src^{CD}$ E283D, W285T | *E. coli* | 261-536 | pET28a, pMCSG7 | N-terminal His6-SUMO tag gets cleaved | This study |
| $Lyn^{myr}$ WT | *S. cerevisiae* | 1-512 | p415 GAL1 | | This study |
| $Lyn^{myr}$ W262T | *S. cerevisiae* | 1-512 | p415 GAL1 | β1/ β 2 resistance cluster mutants | This study |
| $Lyn^{myr}$ E260M | *S. cerevisiae* | 1-512 | p415 GAL1 | β1/ β 2 resistance cluster mutants | This study |
| $Lyn^{myr}$ K275M (kinase dead) | *S. cerevisiae* | 1-512 | p415 GAL1 | β1/ β 2 resistance cluster mutants | This study |

**Table S1: List of constructs and mutations used in manuscript.**

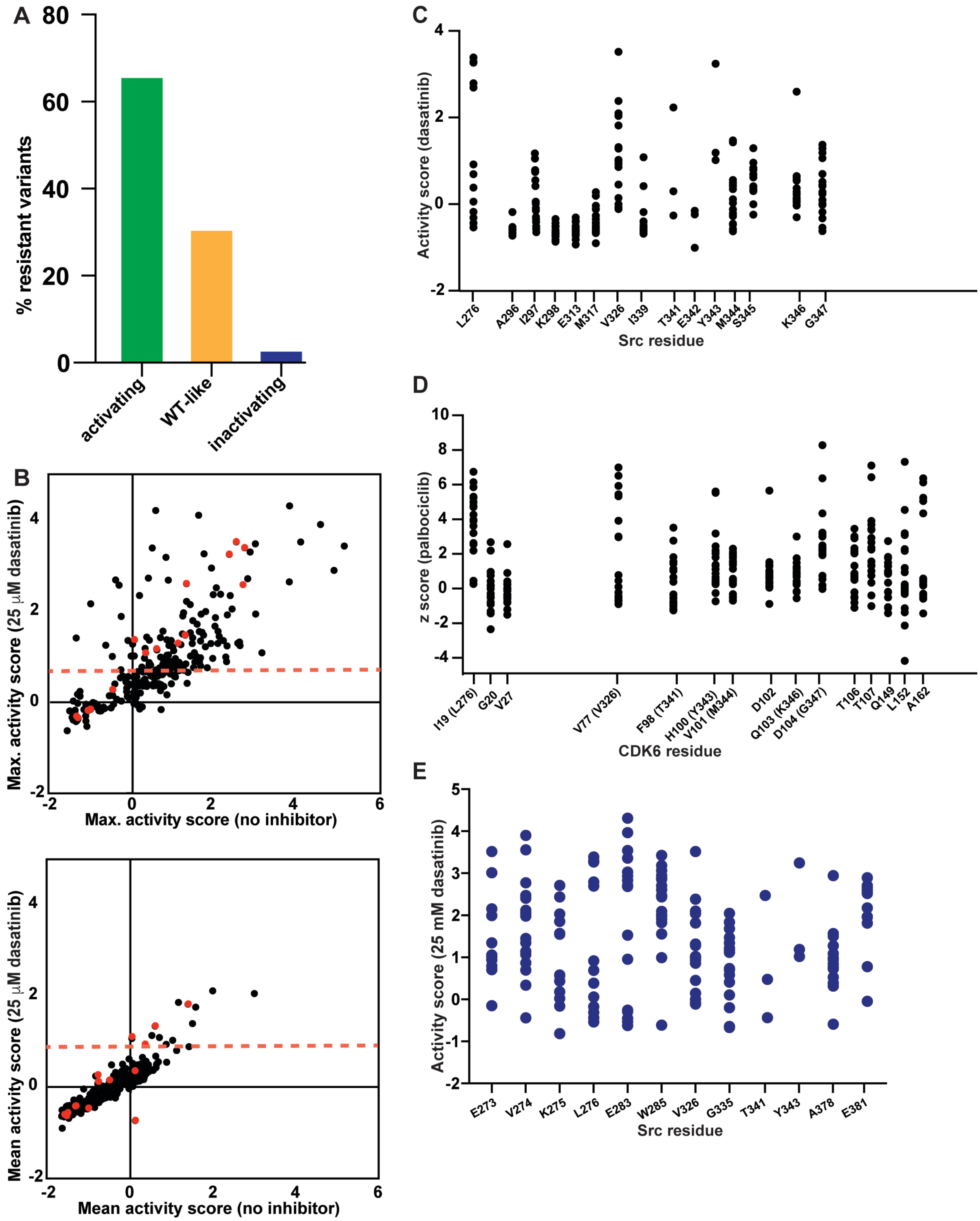


**Figure S2**. **Multiplex analysis of dasatinib resistance in Src.** (A) Percentage of dasatinib-resistant variants that were classified as activating, WT-like, or inactivating in our previous study (Ahler et al., 2019). (B) Correlation plot of maximum activity scores in DMSO-treated (activity scores derived from Ahler et al., 2019) and dasatinib-treated (25 μM) yeast for each residue in Src's catalytic domain. Red dots indicate dasatinib-interacting residues (residues were defined as dasatinib-interacting if they were within 4 Å of dasatinib by LigPlot+ analysis) in the co-crystal structure (PDB ID: 3G5D) of Src bound to dasatinib and black dots indicate residues that do not interact with dasatinib. The red dashed line indicates the activity score value (0.7) we defined as dasatinib resistant ('no inhibitor' data from Ahler et. al. 2019). (C) Scatter plot showing the activity scores measured for each substitution (y-axis, black dots) at dasatinib-interacting residues in Src in the presence of 25 μM dasatinib. (D) Scatter plot showing the z scores for each substitution (y-axis, black dots) at palbociclib-interacting residues in CDK6 (equivalent positions in Src are shown in parentheses). Data derived from Persky et al., 2020. (E) Scatter plot showing the activity scores measured for each substitution (y-axis, blue dots) at the twelve resistance-prone residues in

Src. Residues were defined as resistance-prone if they demonstrated mean activity scores greater than our defined drug-resistance cutoff for an individual mutant in the presence of 25 μM dasatinib. Relates to STAR methods section SFK yeast growth assay.

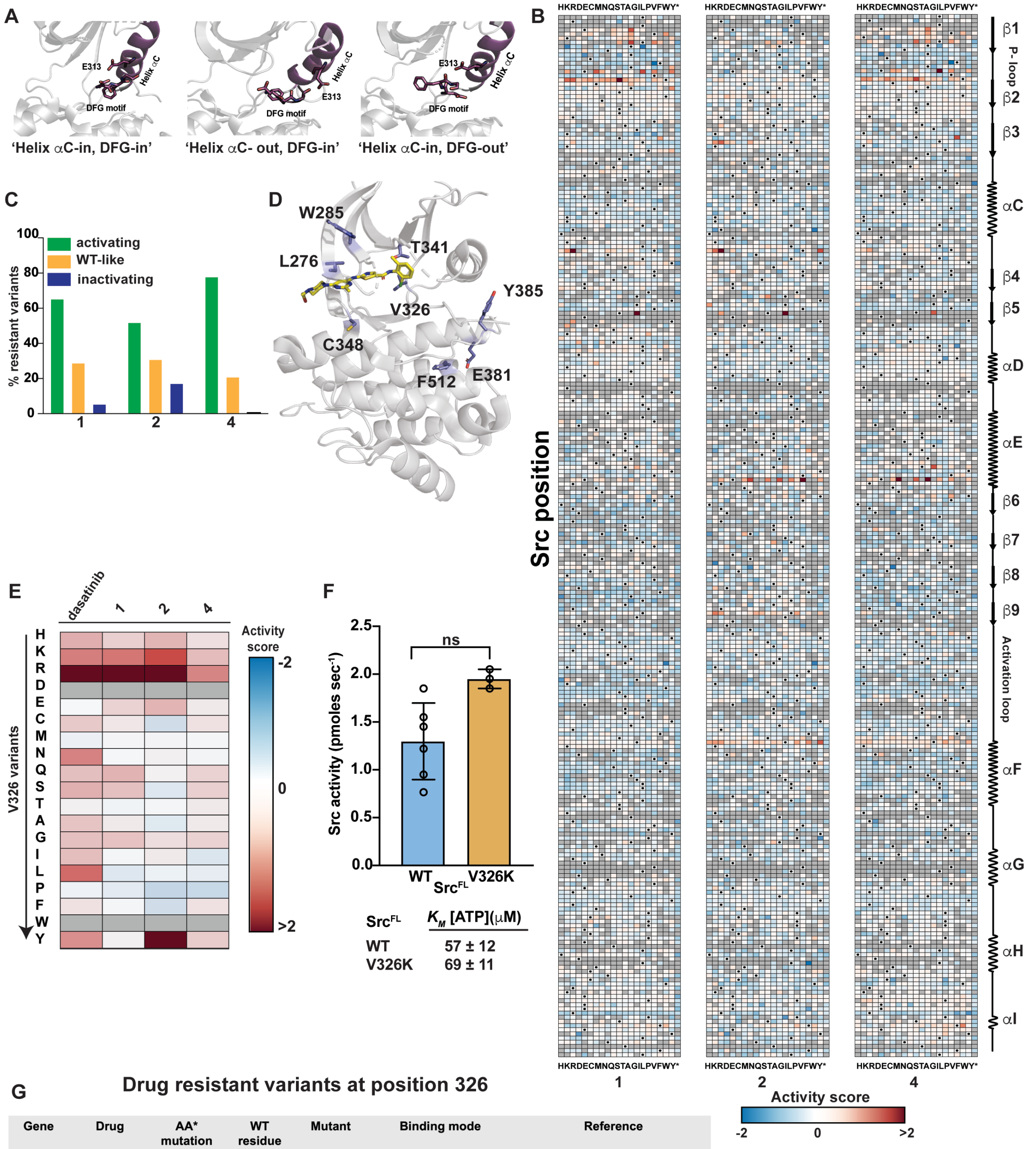


G

**Drug resistant variants at position 326**

| Gene | Drug | AA* mutation | WT residue | Mutant | Binding mode | Reference |
|---|---|---|---|---|---|---|
| KIT | Imatinib | V654A | Valine | Alanine | Helix αc-in, DFG-out | Yongsheng Ma, et.al., 2002 |
| ABL1 | Dasatinib | V299L | Valine | Leucine | Helix αc-in, DFG-in | Elias Jabbour, et.al., 2012 |
| ABL1 | Bosutinib | V299L | Valine | Leucine | Helix αc-in, DFG-out | Elias Jabbour, et.al., 2012 |
| ABL1 | Imatinib | V299L | Valine | Leucine | Helix αc-in, DFG-out | Elias Jabbour, et.al., 2012 |
| ALK | Alectinib | V1180L | Valine | Leucine | Helix αc-in, DFG-out | Ryohei Katayama, et.al., 2014 |

*AA = amino acid

**Figure S3. Parallel analysis of Src's resistance to conformation-selective, ATP-competitive inhibitors.** (A) ATP-binding site conformations stabilized by different conformation-selective inhibitors: (*left*) 'active' with αC helix-in and DFG-in (PDB ID: 3G5D), (*center*) inactive with αC helix-out and DFG-in (PDB ID: 4YBK), (*right*) inactive with αC helix-in and DFG-out (PDB ID: 4YBJ). (B) Sequence-activity score map for all residues in Src's catalytic domain for the $Src^{myr}$ variant library treated with inhibitors **1** (*left*), **2** (*center*), or **4**. Black dots represent the wild-type amino acid and gray tiles indicate missing data. Red indicates higher activity while blue indicates lower activity. Secondary structure and functional motif annotations were obtained from the ProKinO database. (C) Percentage of **1**-resistant (*left*), **2**-resistant (*center*), and **4**-resistant (*right*) mutants that were classified as activating, WT-like, or inactivating in our previous study (Ahler et al., 2019). (D) Crystal structure of Src's catalytic domain (PDB ID: 3G5D) showing the eight residues that demonstrated resistance to inhibitors **1**, **2** and **4**. (E) Activity scores for every substitution at position V326 in the presence of dasatinib or inhibitors **1**, **2**, or **4**. Gray tiles indicate missing data. Red indicates higher activity in presence of inhibitor and blue indicates lower activity in the presence of inhibitor. (F) (*top*) Phosphotransferase activity of purified full-length $Src^{FL}$ with either the WT or V326K sequence, (n=3-6, mean ± SEM).

(bottom) Michaelis-Menten constants ($K_M$ [ATP]) for the $Src^{FL}$ WT and V326K constructs (n=3, mean ± SEM). Data for WT $Src^{FL}$ was reused from Figure 4E. (G) List of drug-resistant mutants at Src position 326 in other tyrosine kinases. Relates to STAR methods section SFK yeast growth assay and Src phosphotransferase activity.

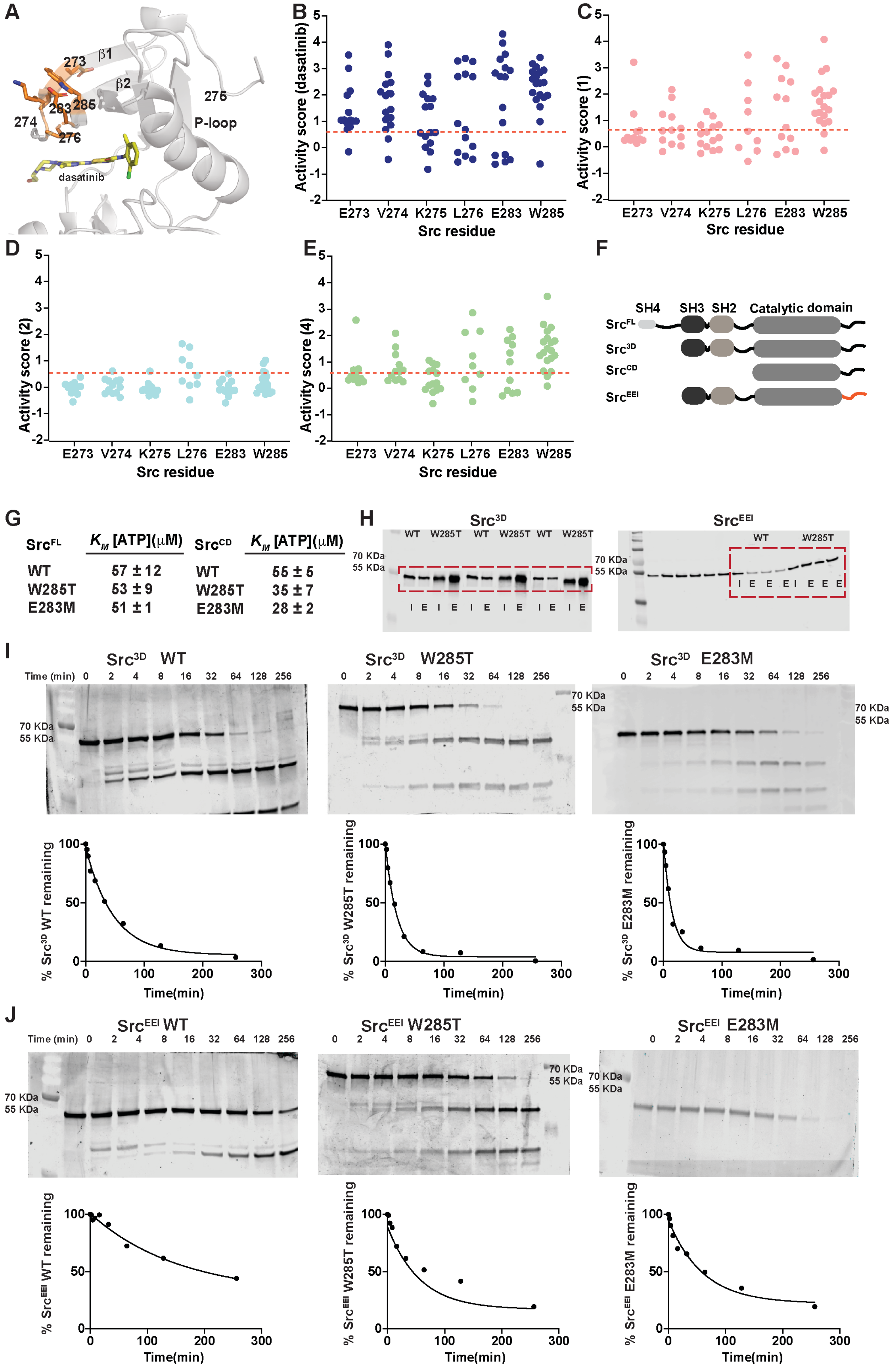
A
273
β1
β2
283
285
274
276
275
P-loop
dasatinib
B
Activity score (dasatinib)
E273 V274 K275 L276 E283 W285
Src residue
C
Activity score (1)
E273 V274 K275 L276 E283 W285
Src residue
D
Activity score (2)
E273 V274 K275 L276 E283 W285
Src residue
E
Activity score (4)
E273 V274 K275 L276 E283 W285
Src residue
F
SH4 SH3 SH2 Catalytic domain
Src^FL
Src^3D
Src^CD
Src^EEI
G
Src^FL  K_M [ATP](μM)
WT 57 ± 12
W285T 53 ± 9
E283M 51 ± 1
Src^CD  K_M [ATP](μM)
WT 55 ± 5
W285T 35 ± 7
E283M 28 ± 2
H
Src^3D
WT W285T WT W285T WT W285T
70 KDa
55 KDa
I E I E I E I E I E I E
Src^EEI
WT W285T
I E E E I E E E
I
Src^3D WT
Src^3D W285T
Src^3D E283M
Time (min) 0 2 4 8 16 32 64 128 256
70 KDa
55 KDa
% Src^3D WT remaining
% Src^3D W285T remaining
% Src^3D E283M remaining
Time(min)
J
Src^EEI WT
Src^EEI W285T
Src^EEI E283M
Time (min) 0 2 4 8 16 32 64 128 256
70 KDa
55 KDa
% Src^EEI WT remaining
% Src^EEI W285T remaining
% Src^EEI E283M remaining
Time(min)

**Figure S4**. **Biochemical characterization of the β1/β2 resistance cluster.** (A) Rotated view of the N-terminal lobe of Src's catalytic domain (PDB ID: 3G5D) bound to dasatinib with the sidechains of all six residues in the β1/β2 resistance cluster shown in orange. (B-E) Scatter plots showing the activity scores for each substitution (y-axis) in the β1/β2 resistance cluster in the presence of dasatinib (B) or inhibitors **1** (C), **2** (D), or **4** (E). The red dashed line indicates the defined activity score (0.7) cutoff for inhibitor resistance. (F) Linear schematic of the various Src constructs used for the biochemical characterization of the β1/β2 resistance cluster. (G) Michaelis-Menten constant ($K_M$ [ATP]) of WT or mutant $Src^{FL}$ and $Src^{CD}$ constructs, (n=3, mean ± SEM; WT $Src^{FL}$ was previously determined in Ahler et al., 2019). (H) Representative Src immunoblots for SH3 pull-down assays performed with WT or W285T $Src^{3D}$ and WT or W285T $Src^{EEI}$. Quantification is shown in **Figure 4G**. (I = input Src construct; E = Src construct retained after SH3 pulldown). Retained Src was quantified by fitting the immunoblot signal intensity to a Src titration standard curve. (I) Representative SYPRO Ruby-stained gel images (*top panels*) and one-phase decay plots (*bottom*) from limited proteolysis experiments (thermolysin) performed with WT (*left*), W285T (*center*), or E283M (*right*) $Src^{3D}$. Quantification is shown in **Figure 4I**. (J) Representative SYPRO Ruby-stained gel images (*top panels*) and one-phase decay plots (*bottom*) from limited proteolysis experiments (thermolysin) performed with WT (*left*), W285T (*center*), or E283M (*right*) $Src^{EEI}$. Quantification is shown in **Figure 4I**. Relates to STAR methods section SFK yeast growth assay, Src phosphotransferase activity, SH3 pull-down assay and limited proteolysis of Src with thermolysin.

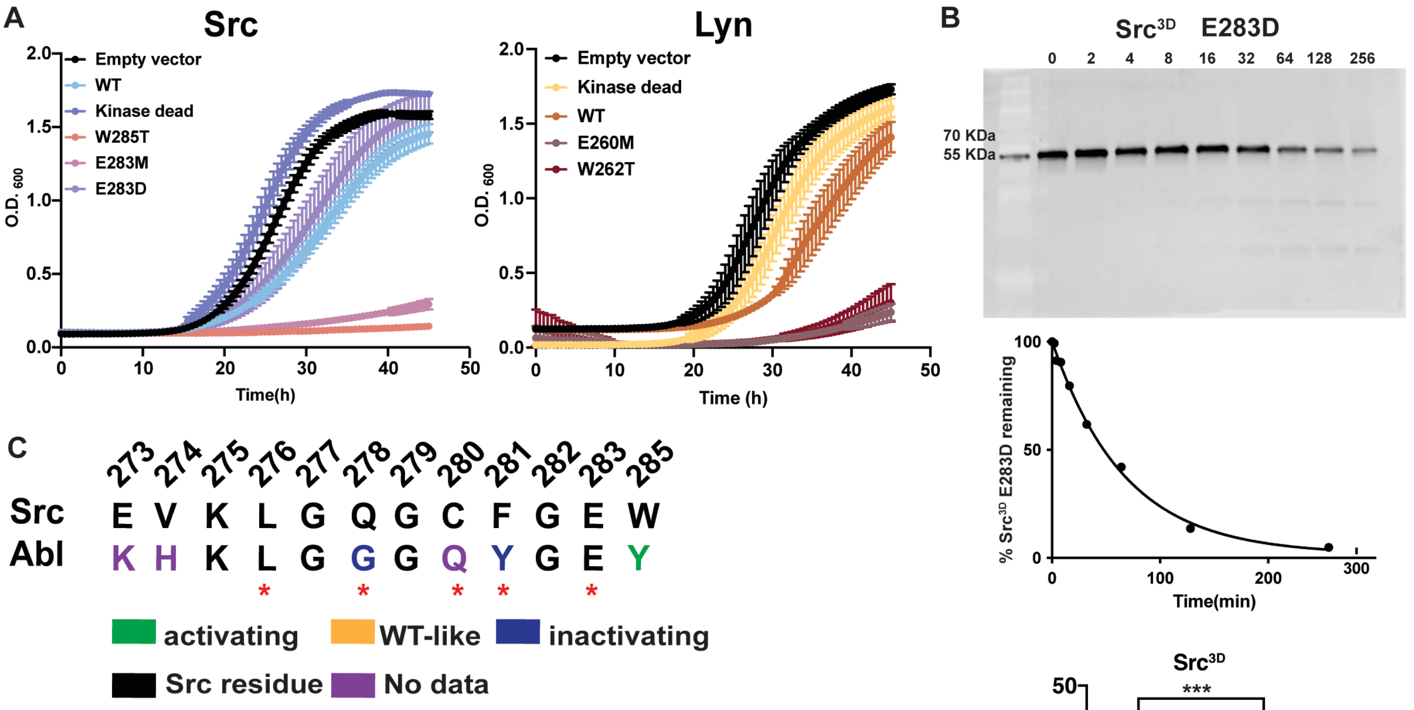


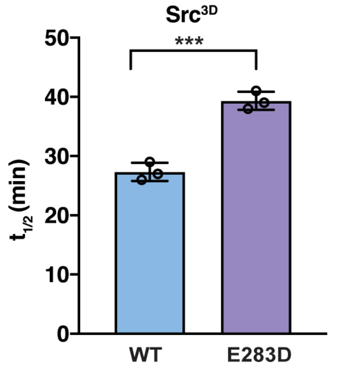


D List of BCR-ABL drug resistant mutations currently in clinic

| Src residue | Location on Src kinase domain | ABL residue | ABL drug resistant mutants |
|---|---|---|---|
| E273 | β1 sheet in N-lobe | K245 | |
| V274 | β1 sheet in N-lobe | H246 | |
| K275 | β1 sheet in N-lobe | K257 | |
| L276 | P-loop | L248 | L248V (#) |
| G277 | P-loop | G249 | |
| Q278 | P-loop | G250 | G250E (+,#,&,%); G250D (#) |
| G279 | P-loop | G251 | G251D (#), G251E (#) |
| C280 | P-loop | Q252 | Q252H (+,#,%); Q252E (#); Q252M (#) |
| F281 | P-loop | Y253 | Y253F(+,#), Y253H (+,#) |
| G282 | P-loop | G254 | |
| E283 | β2 sheet in N-lobe | E255 | E255K(+,#,&); E255V (+,#) |
| V284 | β2 sheet in N-lobe | V256 | |
| W285 | β2 sheet in N-lobe | Y257 | |

Symbol key: Dasatinib (+); Imatinib (#); Bosutinib (&); Nilotinib (%)

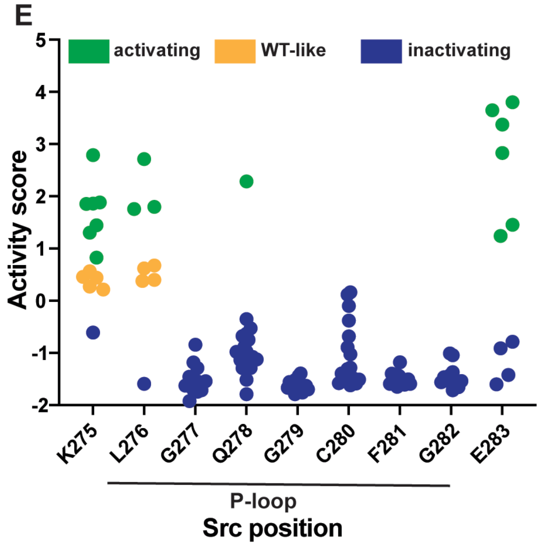


**Figure S6**. **β1/β2 resistance cluster mutations influence the dynamics of Src's P-loop.** (A) (*right*) Individually assessed growth curves for yeast expressing WT, kinase dead, W285T, E283M or E283D variants of $Src^{myr}$ (n=3-6, mean ± SEM). (*left*) Individually assessed growth curves for yeast expressing WT, kinase dead, W262T or E260M variants of $Lyn^{myr}$ (n=3-6, mean ± SEM). (B) Representative SYPRO Ruby-stained gel images (*top panel*) and one-phase decay plot (*center*) from limited proteolysis experiments (thermolysin) performed with WT (*left*), W285T (*center*), or E283M (*right)* $Src^{3D}$. Quantification of triplicate limited proteolysis experiments (*bottom*). (C) Sequence alignment of all the residues spanning the β1/β2 resistance cluster and P-loop in Src with that of the tyrosine kinase Abl. (D) Sites of BCR-Abl inhibitor resistance mutations that have been observed in the clinic. Data obtained from the COSMIC database. (E) Activity scores for every substitution at each position spanning the P-loop of Src in the absence of inhibitors. Activity scores were derived previously (Ahler et al., 2019). Relates to STAR methods section SFK yeast growth assay, and limited proteolysis of Src with thermolysin.
